# Metacell-based differential expression analysis identifies cell type specific temporal gene response programs in COVID-19 patient PBMCs

**DOI:** 10.1101/2023.12.14.571774

**Authors:** Kevin O’Leary, Deyou Zheng

**Affiliations:** Department of Genetics, Albert Einstein College of Medicine, Bronx, NY, USA; Department of Neurology, Albert Einstein College of Medicine, Bronx, NY, USA; Department of Neuroscience, Albert Einstein College of Medicine, Bronx, NY, USA

**Keywords:** scRNA-seq, metacells, SEACells, COVID-19, SARS-CoV2

## Abstract

**Background:** By resolving cellular heterogeneity in a biological sample, single cell RNA sequencing (scRNA-seq) can detect gene expression and its dynamics in different cell types. Its application to time-series samples can thus identify temporal genetic programs active in different cell types, for example, immune cells’ responses to viral infection. However, current scRNA-seq analysis need improvement. Two issues are related to data generation. One is that the number of genes detected in each cell is relatively low especially when currently popular dropseq-based technology is used for analyzing thousands of cells or more. The other is the lack of sufficient replicates (often 1-2) due to high cost of library preparation and sequencing. The third issue lies in the data analysis –-usage of individual cells as independent sampling data points leads to inflated statistics.

**Methods:** To address these issues, we explore a new data analysis framework, specifically whether “metacells” that are carefully constructed to maintain cellular heterogeneity within individual cell types (or clusters) can be used as “replicates” for statistical methods requiring multiple replicates. Toward this, we applied SEACells to a time-series scRNA-seq dataset from peripheral blood mononuclear cells (PBMCs) after SARS-Cov-2 infection to construct metacells, which were then used in maSigPro for quadratic regression to find significantly differentially expressed genes (DEGs) over time, followed by clustering analysis of the expression velocity trends.

**Results:** We found that metacells generated using the SEACells algorithm retained greater between-cell variance and produced more biologically meaningful results compared to metacells generated from random cells. Quadratic regression revealed significant DEGs through time that have been previously annotated in the SARS-CoV2 infection response pathway. It also identified significant genes that have not been annotated in this pathway, which were compared to baseline expression and showed unique expression patterns through time.

**Conclusions:** The results demonstrated that this strategy could overcome the limitation of 1-2 replicates, as it correctly identified the known ISG15 interferon response program in almost all PBMC cell types. Its application further led to the uncovering of additional and more cell type-specific gene expression programs that potentially modulate different levels of host response after infection.

## Background

Single cell RNA sequencing (scRNA-seq) is a powerful tool that can detect distinct gene expression dynamics in different cell types within a sample [1, 2]. One can apply the analysis to time-series samples for the identification of temporal changes in gene expression within each cell type. To do this, a current common practice is to use each cell as a statistical “sample” for determining gene expression change between different time points. Statistically, this is not rigorous because cells in the same biological sample do not really represent independent samples, but have intrinsic correlations [3]. Pseudobulking has been proposed to overcome this, where gene read counts for all cells of a cell type (or cluster) in a biological sample are aggregated. This approach also has an advantage in increasing gene coverage, as relatively low numbers of genes are detected per cell by current scRNA-seq analysis approaches [4]. The strategy, however, highlights the problem of low numbers of replicates in scRNA-seq studies due to the high cost of library preparation and sequencing [5]. In addition, simply aggregating reads in all cells of a type may erase the heterogeneity (or variation) in a cell type (or cluster). In this study, we propose the use of “metacells” to circumnavigate these problems. A metacell represent the transcriptomes of a group of highly similar cells [6]. Multiple methods and algorithms exist to create them [6–8]; however, the single-cell aggregation of cell states (SEACells) algorithm has an advantage in retaining heterogeneity within each cell cluster [9], resulting in metacells representing different states. We thus decided to investigate if the metacells from SEACells can be used as pseudo-replicates (referred as “metareplicates”) in statistical methods that were developed for time-series data from bulk tissues (vs single cells).

Considering the continued importance of understanding the diverse ways in which the immune system responds to severe acute respiratory syndrome coronavirus 2 (SARS-CoV-2), we further decided to test the approach with a time series dataset derived from coronavirus disease 2019 (COVID-19) patients following symptom onset [10].

SARS⍰CoV⍰2, the strain of coronavirus responsible for the coronavirus disease 2019 (COVID-19) pandemic [11, 12], continues to infect hundreds of thousands of people around the globe. To date, almost 7 million confirmed deaths have been recorded as a consequence of SARS-CoV-2 infection [13]. The desire to understand the mechanisms behind SARS-CoV-2 infection and host defense, especially as it relates to its transmissibility [14, 15] and severity [16], has prompted a vast amount of research in the field of immunology and beyond [17, 18]. One of many topics of interest concerns gene programs within cell types that respond to SARS-CoV-2, specifically peripheral blood mononuclear cells (PBMCs), which are any round nucleus containing blood cells such as dendritic cells, lymphocytes, natural killer cells (NKs), or monocytes [19]. Because PBMCs are responsible for responding to and eliminating viral infections such as SARS-CoV-2, it is important to understand the transcriptomic basis of this process.

Researchers have compared gene expression in PBMC cell types between COVID-19 patients and controls using bulk RNA sequencing [20]. Others have implemented scRNA-seq [21, 22], which provides greater resolution at the cellular level, especially as it relates to deducing cell type-specific responses to SARS-CoV-2 infection. Some have even performed time series scRNA-seq analysis of COVID-19 progression. While these studies have provided valuable information relating to cell type-specific changes in expression through time, they were limited by the issues of small replicates as discussed above. For example, some time points in the PBMC scRNA-seq data that we planned to analyze have only 1 replicate. Consequently, the authors had to bin samples of different time points to increase statistical power [10]; so did in other studies [20, 23, 24]. In addition to this computational difference, the scope and focus of our current study is also different from the original report [10], e.g., the original authors focused on the response difference between COVID-19 infection and flu and did not study the velocity of the expression changes. The authors of SEACells also studied SARS-CoV-2 gene responses in PBMCs with a different dataset [21], but focused on CD4 T cells and only analyzed a few metacells that differed based on dominance of certain time points [9]. This differs from our study in that we analyzed metacells representing 10 discrete points in time and in many PBMC cell types.

In short, using the SEACells alogorithm, we created metacells that retained hetergeneity within each cell type and used them as metareplicates. This resulted in up to 12 replicates for some time points and thus provided the statistical power necessary to resolve signficiant changes in expression through time. With that, we performed strict statistical analysis through a greater number of time points than any other COVID-19 time-series scRNA-seq study to date. To accomplish this, we subset all cells based on time since symptom onset and then used the SEACells algorithm to create metacells. maSigPro [25] was used for quadratic regression to find significantly differentially exprssed genes (DEGs) through time, due to its robust statistical base, its flexibility with defining degrees of regression, and widespread use for time series analysis. Additionally, quadratic regression was used because we did not want to capture cyclical variation, rather we hoped to find broader changes in expression through four weeks of COVID-19 symptoms. We further classified all DEGs by expression velocity trend based on fitted expression curves and their dynamic derivates. With this approach, we identified*ISG15* as a DEG through time when PBMC cell types were analyzed together. When cell types were analyzed independently, however, we found many immune system-related DEGs, which enabled us to expand upon previous reports of certain gene programs and their relevence to SARS-CoV-2 immune response.

## Methods

### Metacell Creation

The COVID-19 scRNA-seq dataset was obtained from a previous study that performed time series analysis on PBMCs from five SARS-CoV-2 infected patients [10]. The date of symptom onset and sample collection was recorded for each patient. Since we did not intend to group patients by disease stage, we simply classified each collected sample by the number of days after symptom onset. Samples from influenza patients were excluded from time series analysis, as were controls, since they were not collected continuously through time. However, we included the normalized expression of three healthy controls as baseline values for comparative purposes.

For SEACells, the number of metacells was determined based on the software authors’ suggestion of 1 metacell per 75 single cells [9]. We rounded to the nearest 10 to enable the creation of more metacells for time points with fewer total cells. The assignment of individual cells to metacells was determined using the SEACells algorithm in Python. We applied SEACells to samples of each time point independently. For each of the 10 time points, the input consisted of an Anndata object containing normalized counts from the n most highly variable genes (2000 for our dataset), cell cluster/type assignments (as previously determined by the original author [10]), and a low dimensional representation of the data. Subsequent steps for metacell creation were outlined in the SEACells manuscript [9] and in **Figure 1**. The expression of each gene for a given metacell was determined by averaging the normalized counts of the cells that were assigned to it **(Figure 1A-B)**. Each metacell was ascribed a cell type based on whichever cell type was most prominent amongst the assigned individual cells. For example, if most cells assigned to a metacell were plasma cells, the metacell would be called a plasma metacell. The percentage of cells comprising the metacell that were of the assigned cell type was referred to as its “purity.” We call metacells created using the SEACells algorithm “sMetacells.” To obtain metacells composed of random individual cells by cell type, which we call “rMetacells”, we subset the same filtered PBMC dataset by time. We then subset by cell type and took the average normalized expression of 20 randomly selected cells to create an rMetacell. While we intended to ues more cells to create metacells that were as comparable to sMetacells as possible, several cell typeshad less than 75 cells for a particular time point, so we decreased our threshold to maximize metacell assignments. The SEACells algorithm was not confined to this issue due to its ability to assign varying numbers of cells to each metacell based on nearest neighbor determinations.

**Figure 1.**
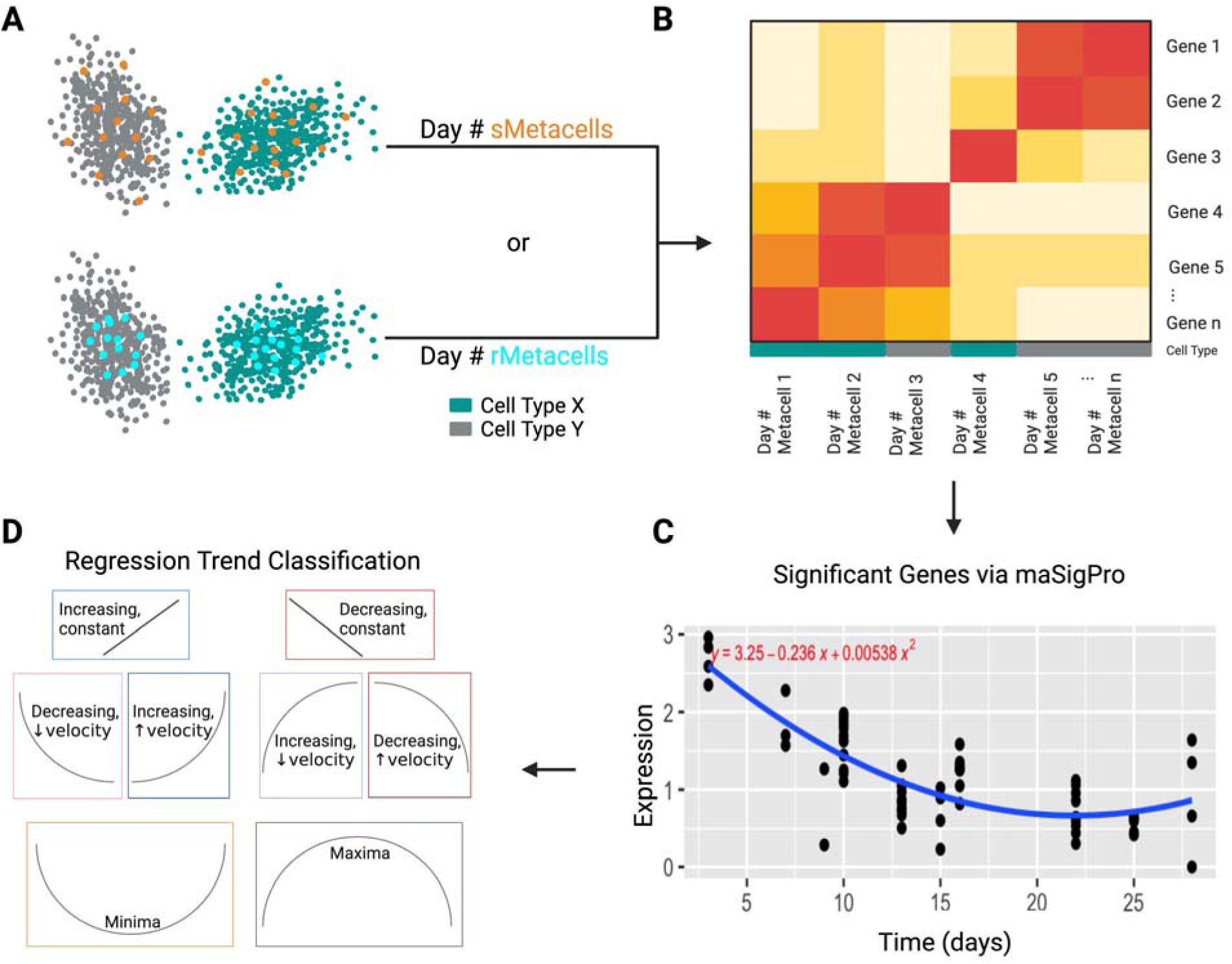
(previous page): Summary of metacell generation and usage. **A)**An example of metacells generated using the SEACells algorithm (sMetacells) and random single cells (rMetacells) for a time point. An example of the distribution of sMetacells (orange dots) and rMetacells (blue) are shown overlaying all cells of a particular type (either grey or blue). sMetacells, given their propensity to distribute over the full cell type space, are more spread out while rMetacells depend on random assignment of cells and therefore have a higher probability of occupying the space with greatest cell density. **B)** The gene expression of each metacell was computed from the average of the normalized expression amongst all single cells assigned to it. **C)** After the generation of metacells for each time point, quadratic regression was performed for each gene. An example of a significantly changing gene is shown here. **D)** One of eight expression trends was assigned to each DEG. For the example in C, the trend would be “Decreasing, ↓ velocity” for decreasing expression with decreasing

### maSigPro and Trend Determination

After the creation of metacells (by SEACells or randomly) for each time point, maSigPro was used to find DEGs through time. maSigPro utilizes regression ANOVA followed by a variable selection procedure [25]. Quadratic regression was used since we expected the change in gene expression to follow one of eight general trends, as described in **Figure 1C-D**. For each cell type and gene, a quadratic equation was generated to represent expression through time. Only genes with false discovery rate (FDR) less than 0.05 and R^2^ value greater than 0.5 were considered statistically significant and retained. An F statistic-associated p-value was produced for each coefficient A, B, and C in equation 1.

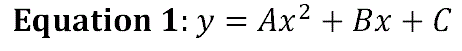

The trends with p < 0.05 for coefficient A were determined based on the shape of their fitted curve. If the absolute value of the slope of the line tangent to the expression vs time curve (the expression velocity) decreased through time, we called this decreasing velocity, denoted “↓ velocity” in figures. If the absolute value of the slope of the tangent line increased, this was referred to as increasing velocity, or “↑ velocity.” We combined these terms with the overall trend of increasing or decreasing expression. For example, if the expression of a gene was decreasing through time, was not linear, and showed decreasing velocity, we would call this decreasing expression with decreasing expression velocity or “Decreasing, ↓ velocity” for short. If p > 0.05 for coefficient A, we considered this to be linear and the direction of the curve dictated whether it was considered increasing or decreasing. Increasing linear expression is synonymous with “Increasing, constant” while decreasing linear expression is synonymous with “Decreasing, constant” where constant refers to the expression velocity. If the average expression for the first time point and the last time point were both less than each of the time points between them, this was considered “Maxima”. If greater, this was “Minima”.

It is important to note that DEGs from dendritic cells (DCs), megakaryocytes, monocytes, cycling plasma, and stem cells were eliminated from further analysis due to low cell numbers (less than 500 in total across all time points), which led to numbers of metacells too low for robust statistical analysis, because performing quadratic regression would lead to overfitting for these cell types. Additionally, due to low metacell counts for the first three time points in memory B cells, we eliminated days 3, 7, and 9 metacells for trend determination of this cell type due to skewing toward early time point outliers. For all other cell types, all ten time points (days 3, 7, 9, 10, 13, 15, 16, 22, 25, and 28) were included for trend determination.

### Other Bioinformatics Databases and Tools

For classification of the functions of the gene products (i.e., proteins), we used the DAVID Gene Function Annotation Tool [26, 27] and further grouped selected terms into broader function categories, such as transferases, proteases, immunoglobulin-related, and interferon-related. The KEGG [28] COVID-19 pathway was used to define known SARS-CoV-2-related genes. Although the KEGG pathway is based on SARS-CoV-2 entry into type 2 pneumocytes, we generalized this response to the cascade of events that follow uptake of the virus by PBMCs to further narrow our search for novel expression responses. We base this generalization on the finding that cell-intrinsic innate immune responses are triggered in PBMCs following exposure to SARS-CoV-2 [29]. The STRING Database [30] was used for network analysis to connect our DEGs to known COVID-19-related genes. To find significantly enriched gene ontology (GO) terms from inputted DEGs, we used geneontology.org [31, 32], set the annotation dataset to “PANTHER GO-slim biological processes”, and used the entire human genome as background. Figures were edited using biorender.com.

## Results

### Finding DEGs through time with pseudobulking method

To characterize the dynamics of cell type gene programs in the PBMCs in response to SARS-CoV-2 infection, we first applied a pseudobulking approach by aggregating scRNA-seq reads for individual genes, for either all PBMC cells or each of the cell types, for each sample. The samples and scRNA-seq data were collected at 10 time points representing post symptom onset days, from day 3 to day 28 by Zhu et al, as described previously [10]. This generated timeseries pseudobulk RNA-seq data with 1 to 3 replicates, which were then used to identify genes exhibiting significant expression changes along the post infection period by maSigPro. The regression ANOVA analysis did not find any DEGs when PBMCs were not separated into cell types but found a few DEGs for some cell types (1 for T cells and 3 for plasma cells) (**Figure S1**). However, most of the DEGs exhibited the same expression trend, suggesting model overfitting due to outliers and low replicates.

### Characterizing DEGs through time using metacells as replicates

We reasoned that using metacells to construct computational replicates (referred as “metareplicates”) may allow us to mitigate false positives and overfitting in the pseudobulk approach. To test this, we generated metacells from the scRNA-seq data for samples in each of the 10 time points independently using two different methods: SEACells and random selection (see Methods). The resultant metacells were referred as “sMetacells” and “rMetacells”, respectively. Given that the SEACells algorithm retains heterogeneity within specific cell types, we expected that its metacells would introduce variation within individual time points and lead to fewer DEGs through time and be less prone to overfitting. We therefore compared the numbers of DEGs determined for these two methods **T**( **able 1**). We excluded all cell types with fewer than 500 total cells to avoid more extreme cases of overfitting for both methods since fewer cells would lead to fewer metacells (and thus few replicates). After that, the average number of replicates per time point for each cell type using the SEACells algorithm was 3.3. For rMetacells, three replicates were created. The average number of cells assigned to each metacell was 31.8 for sMetacells and predetermined to be 20 for rMetacells.

For each metacell type, we determined the standard deviation (SD) of each gene’s expression for each time point and used these values to calculate the mean SD (mSD) for all genes. Thus, we obtained mSD values for each time point and cell type for either rMetacells or sMetacells. sMetacells showed greater mSD, and therefore greater variance, for 72 out of 100 individual time points across cell types. Additionally, if these mSDs were further averaged among all time points and cell types, sMetacells still showed greater mSD (0.065) than rMetacells (0.041), a difference that was statistically significant (p = 2.27e-7, t-test). These results are summarized in**Table S1**. Overall, this indicates that sMetacells provide bigger variances among metareplicates than rMetacells.

A more important question is how the variances provided by metacells match the true biological variances. Since at individual data points there were insufficient biological replicates to provide good estimate of sample variances, we decided to combine cells from all time points and computed gene SDs for each of the cell types, with pseudobulking, rMetaCell, or sMetacell methods. The result indicated that the gene variances from sMetaCells were very close to those from pseudobulking and significant larger than those from rMetaCells (**Figure S2**), further supporting that sMetacells could be used as replicates. Interestingly, the average number of genes per metacell was also higher for sMetacells (8,440) than rMetacells (5,930) (p = 2.2e-16, t-test) (**Table 1**).

**Table 1:** Comparison of metareplicates from sMetacells and rMetacells. A, Replicates per time point, # cells per metacell, average variance, and average # of genes detected. B, The number of DEGs through time using quadratic regression at FDR < 0.05 and R^2^ > 0.5.

|  | Metacell Method | sMetacells | rMetacells |
| --- | --- | --- | --- |
| A<br>(Summary<br>of<br>Metacells) | Metareplicates Per Time Point | 3.3 | 3 |
|  | Avg # of Cells Assigned to Metacell | 31.8 | 20 |
|  | Avg Variance Across Gene | 0.041 | 0.0097 |
|  | Avg # Genes in Metacells | 8440 | 5930 |
| B (DEGs) | All PBMCs without separating to cell types | 1 | 10 |
|  | Cytotoxic CD8 T cells | 38 | 31 |
|  | Naïve T cells | 19 | 22 |
|  | NKs | 25 | 43 |
|  | Activated CD4 T cells | 74 | 64 |
|  | Naïve B | 33 | 91 |
|  | Plasma | 7 | 120 |
|  | Memory B | 9 | 57 |
|  | XCL+ NKs | 15 | 79 |
|  | MAIT | 53 | 49 |
|  | Cycling T cells | 68 | 633 |
|  | Total DEGs | 342 | 1199 |

Performing quadratic regression yielded more DEGs using rMetacells than sMetacells, likely due to a higher degree of overfitting due to less variation across metareplicates **T**( **able 1B**). However, the difference between the total number of DEGs found using sMetacells vs rMetacells was not statistically significant. Regardless, for cycling T cells, over 600 DEGs were detected for the rMetacell method compared to 68 using sMetacells. Of all the DEGs from the two methods, 49 were the same, leaving116 and 984 unique to the sMetacell and rMetacell methods, respectively. To better undrestand the difference, we performed gene ontology (GO) enrichment analysis using all the DEGs identified fromat least one of the cell types (FDR < 0.05) (Figure 2). The results showed that the DEGs from the sMetacell method, despite fewer in number, were actually enriched with more significant GO terms, particularly those related to immune response. Additionally, for “defense response to virus” and “response to virus” terms, which were significant using DEGs from both methods, the fold enrichment scores were greater and FDR values were lower from results produced by sMetacells. This indicates that DEGs from sMetacells are more biologically relevant and less likely from statistical noise (i.e., false positives), e.g. overfitting due to underestimated variance by rMetacells. We therefore consider the metacells from the SEACells algorithm to be more appropriate metareplicates and discuss results from this method further in more details.

**Figure 2:**
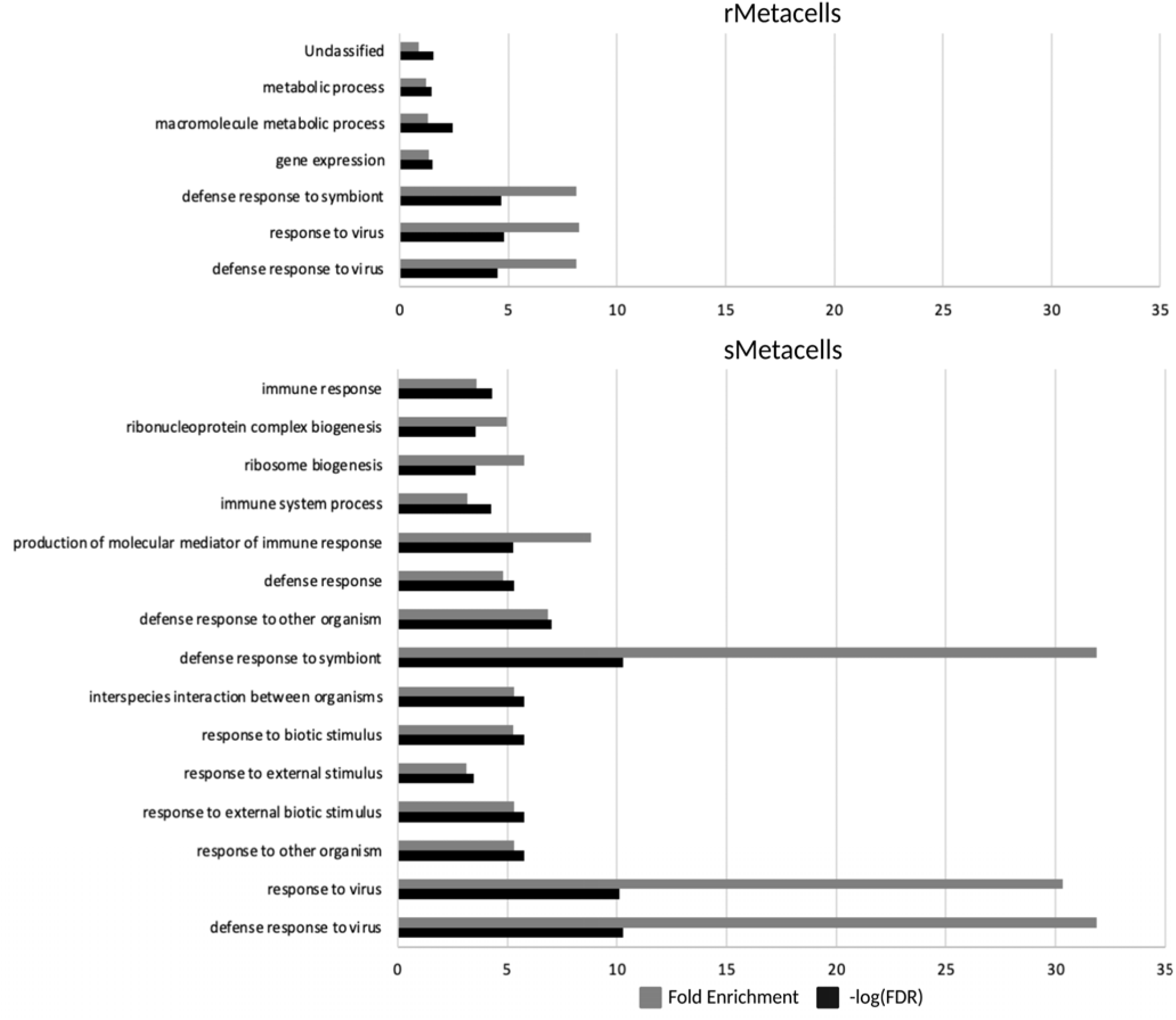
SEACells-derived DEGs show stronger enrichment of biologically relevant pathways than those derived from random metacells. PANTHER GO-slim biological processes annotation data set was used to find enriched terms amongst DEGs through time from rMetacells and sMetacells.

**Table S2** summarizes the number of samples, cells, and metacells for each time point using the SEACells algorithm. The cell identity of each metacell was assigned to the most abundant cell type among the individual single cells contributing to the metacell, using the metadata provided by Zhu et al. **Figure 3A** shows a UMAP representing the 25,775 cells from the COVID-19 patients and their assignment to one of the fifteen cell types. The SEACells algorithm performed exceptionally well in creating metacells that encompass the entirety of the cell type and state space for each time point **(Figure 3B**). As expected, sMetacells had significantly higher numbers of genes detected (8,840 on average) compared to single cells (814 on average) (**Figure 3C**). The proportion of cells in each sMetacell that were from the same cell type were very high, indicating high sMetacell purity, with the average purity scores reaching 90% or higher (**Figure 3D**).

**Figure 3.**
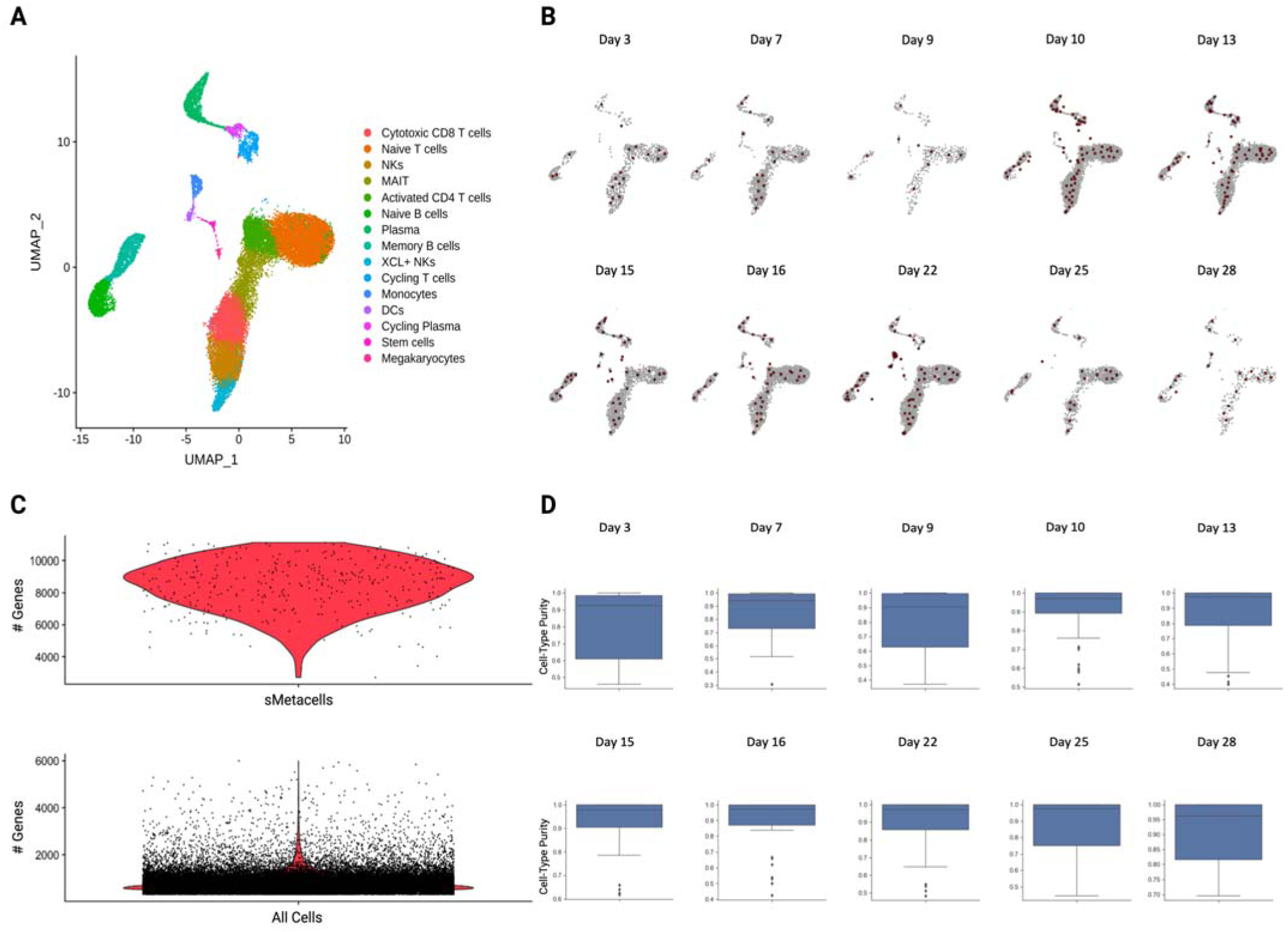
(next page): Summary of sMetacell output. **A**) UMAP of 25,775 cells colored by cell type. **B)** Metacell distribution across cell type space for each time point. Metacells are red while single cells are grey**C**. ) Violin plot of the number of genes detected for SEACell-generated metacells (top) compared to all single cells (bottom). **D**) Box plots showing metacell purity for each day. The average purity for each metacell was over 90% for all time points.

After eliminating cell types with low cell number and with too few sMetacells to be used as metareplicates in maSigPro analysis, we identified 165 unique DEGs through time with an R^2^ > 0.5 and FDR < 0.05, with some DEGs found in more than one cell type (Table S3). We grouped the DEGs based on their functions and the cell type in which they were identified. Within each cell type, genes were futrher grouped according to the expression trends along the times (Figures 1, 4). The trends for all significant DEGs through time by cell type can be found in Figure S3. The results showed that activated CD4 T cells contained the greatest number of DEGs, followed by cytotoxic CD8 T cells, naïve B cells, natural killer cells (NKs), XCL+ NKs, Naïve T cells, Memory B cells, and Plasma cells, respectively (Figure 4B). As mentioned previously, low overall numbers of monocytes, DCs, cycling plasma, stem cells, and megakaryocytes led to low metacell numbers for these cell types, so they were eliminated from furhter analysis. Additionally, upon visual inspection of clustered DEG trends for MAITs and cycling T cells (**Figure S4**), we found that a large group of genes had zero expression but were influenced by an early time point outlier, which led to overfitting. We therefore also eliminated these cell types from further analysis.

**Figure 4.**
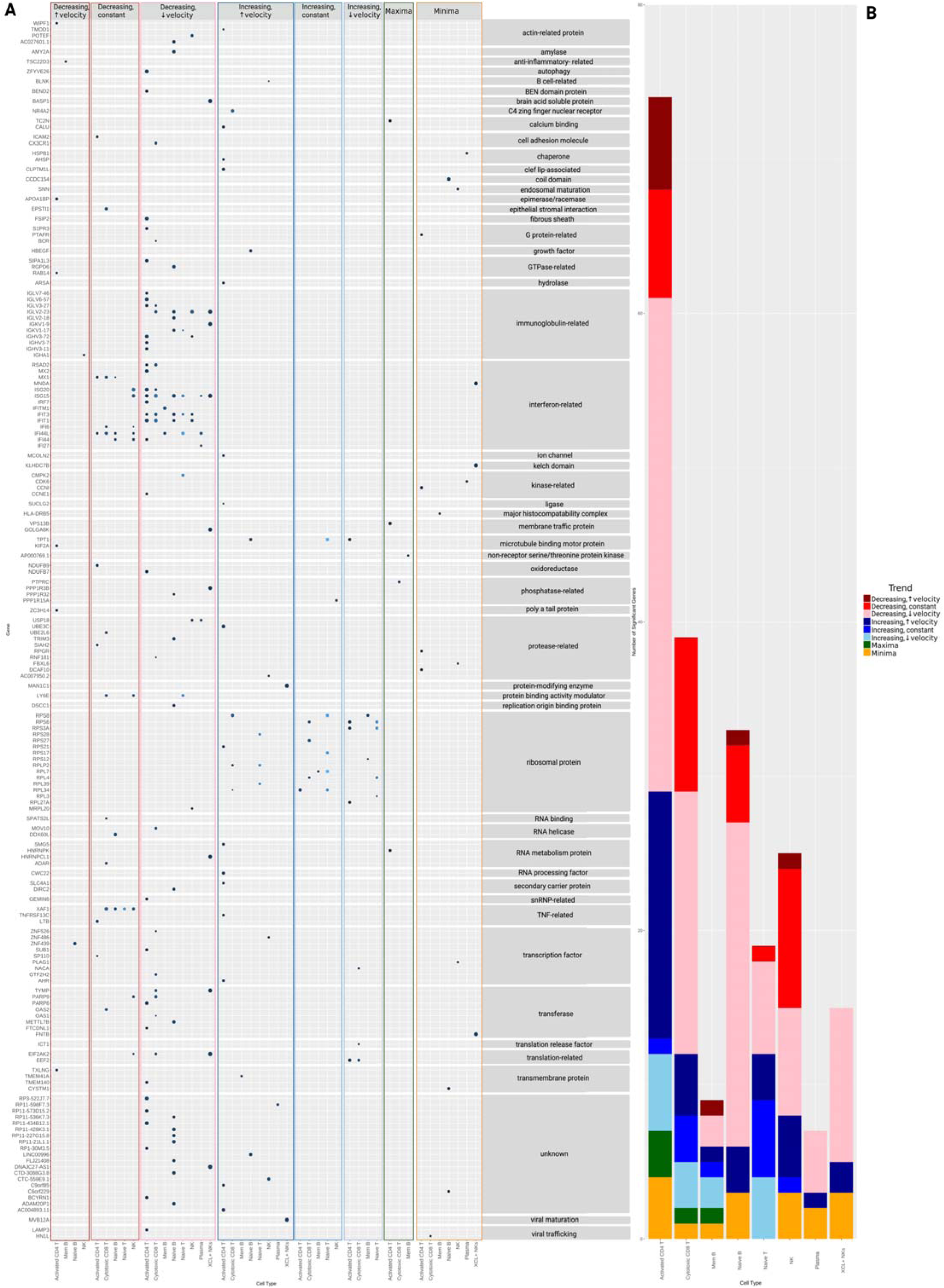
(next page): Summary of significant DEGs and expression trends by cell type. **A**) Dot plot of all significant DEGs through time by trend type, protein class, and sMetacell type. A lighter blue dot corresponds to a lower p-value while a larger dot represents a larger R^2^. **B)** Summary of expression trends by metacell type. The y-axis corresponds to the frequency of significant DEGs through time for each cell type that correspond to a given trend pattern. Red shades represent overall decreasing expression through time, blue shades are increasing, green is maxima (increasing then decreasing) and orange is minima (decreasing then increasing).

At the level of general functional categories, the largest proportion of the DEGs were related to ribosome (16), followed by interferon (14), immunoglobulin (11), protease (10), transcription factor (9) and transferase (8). The functions for 20 of the DEGs was unknown. These results are summarized as a dot plot in **Figure 4A**. 15 of the 16 ribosome-related genes showed at least one of the three increasing gexpression patterns through time depending on cell type while 1/16 (*MRPL20*) showed decreasing expression with decreasing velocity in NKs. 13/14 interferon-related genes showed either a linear decreasing expression pattern or decreasing expression with decreasing expression velocity in a variety of cell types while 1/14 (*MNDA*) showed a “minima” trend in XCL+ NKs. 10/11 immunoglobulin-related genes showed decreasing expression with decreasing velocity through time while 1/11 *I*(*GHA1*) showed a linear decreasing trend in NKs. Protease and transcription factor-related genes fit into a variety of increasing, decreasing, and minima trends depending on cell type. 7/8 transferase genes showed either linear decreasing expression or decreasing expression with decreasing velocity while 1/8 *F*( *NTB*) showed a “minima” trend in XCL+ NKs.

### Connecting many DEGs to genes previously implicated to SARS-CoV-2 response

Next, we asked how the 165 DEGs are related to genes previously found to be involved in the COVID-19 pathway, based on KEGG. We input the protein names corresponding to these 165 genes into the StringDB to determine protein associations and colored the nodes by whether they are in the COVID-19 KEGG pathway (**Figure 5A**). 89 of the 165 genes were determined to have a protein product that interacted with at least another protein from the input. Among these 89, 23 were previously annotated as being part of the KEGG COVID-19 pathway. We also colored nodes by the protein’s affiliation with significant biological processes that capture the three main clusters of connected proteins (**Figure 5B**). Actual cluster identification prior to overlay with biological process identifiers can be found in **Figure S5.** The analysis showed that the DEGs were significantly enriched for functions related to Translation (FDR = 1.75e-09), Cell Surface Receptor Signaling (FDR = 0.00023), and Type I Interferon Signaling (FDR = 8.45e-14). The proteins comprising the Translation cluster are ICT1, MRPL20, NACA, RPL7, RPL27A, RPL34, RPS17, RPLP2, RPL39, EEF2, RPS8, RPL3, RPS27, RPS12, RPS28, RPL4, RPS21, RPS6, RPS3A, RPS27, and EIF2AK2. Among these, NACA, ICT1, MRPL20, and EEF2 were not previously annotated in the COVID-19 KEGG pathway. All proteins belonging to the Type I Interferon Signaling group overlapped with the Cell Surface Receptor Signaling group. These proteins include ADAR, IFI27, ISG20, ISG15, XAF1, MX2, RSAD2, IFIT3, IFITM1, IFI6, OAS1, OAS2, IFIT1, MX1, and RF7. Among these, IFI27, ISG20, XAF1, RSAD2, IFIT3, IFI6, IFITM1, IFIT1, and IRF7 were not previously annotated in the COVID-19 KEGG pathway. Proteins annotated in only the Cell Surface Receptor Signaling group were HSPB1, CCNE1, CDK6, MOV10, LY6E, NR4A2, PTPRC, ICAM2, MNDA, BCR, BLNK, IGHV3-11, S1PR3, HBEGF, CX3CR1, HLA-DRB5, LTB, and TNFRSF13C. Among these genes, none except HBEGF were previously annotated in the COVID-19 KEGG pathway. The results indicate that DEGs from our analysis likely have important roles in modulating immune responses.

**Figure 5:**
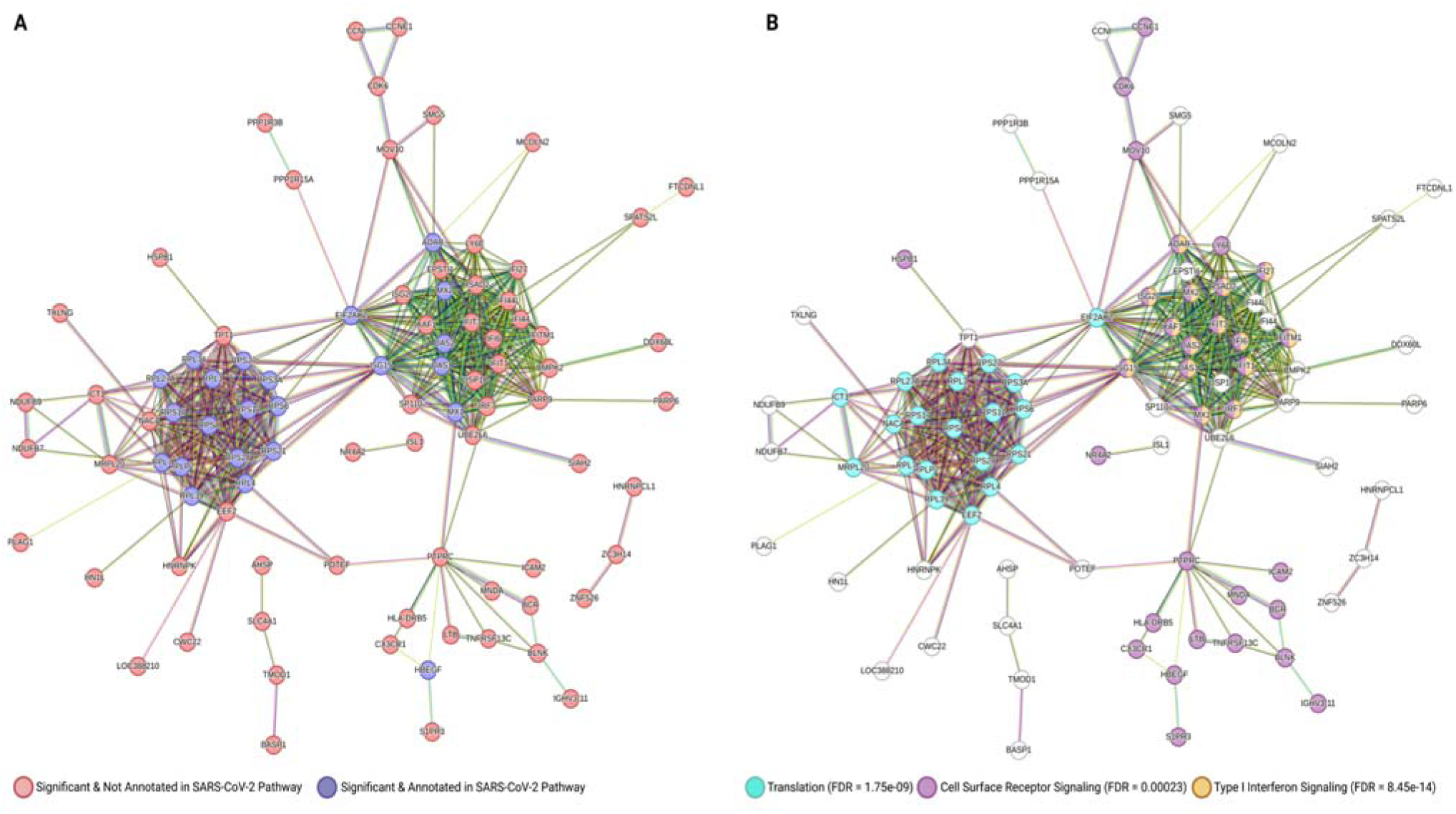
STRING protein interaction results. **A**) STRING network colored by annotated vs unannotated KEGG COVID-19 pathway-related protein products. Red represents protein products from genes that are not annotated in the KEGG COVID-19 pathway. Dark blue represents those that are already annotated in this pathway. **B**) STRING network colored by Biological Process GO Terms. GO terms were selected based on their ability to encompass 3 main clusters. Turquois represents Translation, purple represents Cell Surface Receptor Signaling, and yellow/orange represents Type I Interferon Signaling.

### Detailed description of DEGs newly implicated to SARS-CoV-2 response

As these DEGs changed expression post-infection, we wondered if their day 3 expression would be significantly different between infected PBMCs and controls. We also wondered whether at day 28 their expression would return to the baseline (**Figure 6**). To illustrate this, we plotted the expression of DEGs associated with one of three significant GO biological process terms but not in the KEGG COVID-19 pathways, i.e. genes that are not yet well described in COVID-19 literature (**Figure 5**). We compared the expression at day 3 between COVID-19 infected cells and healthy controls and found that 31 of the DEGs exhibited a significant difference (t-test) except those showing a “minima” or “maxima” trend, as expected. Given the low number of metacells for day 28 data, we did not perform this test but the expression of almost all DEGs were back to the baseline levels (**Figure 6**).

**Figure 6.**
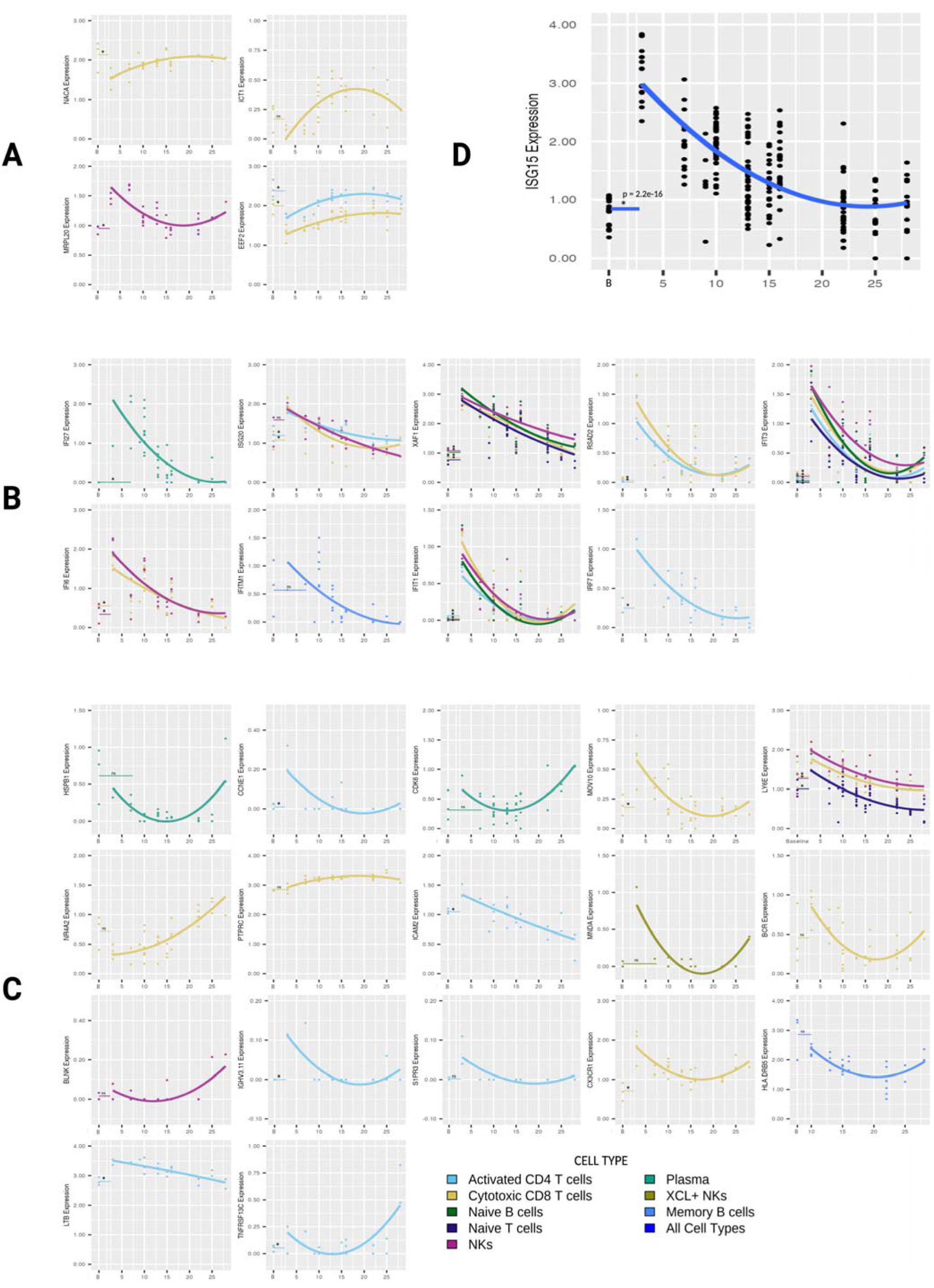
(previous page): Expression vs time plots and trendlines for selected significant genes. **A**) Expression through time for “Translation” genes that do not overlap with the KEGG COVID-19 pathway. **B)** Expression through time for “Type I Interferon Signaling” genes that do not overlap with the KEGG COVID-19 pathway. **C)** Expression through time for “Cell Surface Receptor Signaling” genes that do not overlap with the KEGG COVID-19 pathway.**A-C)** Only cell types whose expression of a gene was deemed significant through time were plotted.**D)** ISG15 expression through time for all cell types together. **A-D)** All baseline values were compared to day 3 (or day 3 and day 7 for plasma cells) via two-sided, unpaired t tests with equal variance. For each cell type, lines matching their associated color were drawn to represent the baseline average. Significant differences between baseline and day 3 expression were denoted by an asterisk. “B” at the x-axis represents expression in healthy controls (baseline).

For genes whose protein products are related to translation (**Figure 6A**), *NACA* showed increasing expression with decreasing expression velocity in cytotoxic CD8 T cells, as did*ICT1* and *EEF2*. *EEF2* also demonstrated this trend in activated CD4 T cells.*MRPL20* exhibited decreasing expression with decreasing expression velocity in NKs. Expression at 28 days closely resembled that of baseline expression for these four genes.

Among genes whose protein products are related to cell surface receptor signaling (**Figure 6B**), *IFI27* expression decreased through time with decreasing expression velocity in plasma cells.*ISG20* expression decreased through time with decreasing velocity in activated CD4 T cells and cytotoxic CD8 T cells. Expression of *ISG20* showed a linear decrease through time in NKs. Naïve T cells, naïve B cells, cytotoxic CD8 T cells, and NKs exhibited a linear decrease in*XAF1* expression. *RSAD2* showed decreasing expression with decreasing velocity in activated CD4 T cells and cytotoxic CD8 T cells while*IFIT3* exhibited the same trend in naïve T, naïve B, activated CD4 T cells, cytotoxic CD8 T cells, and NKs*I*.*FI6* showed a linear decrease in expression through time for cytotoxic CD8 T cells and NKs.*IFITM1* expression decreased through time with decreasing expression velocity in memory B cells.*IFIT1* showed the same trend in naïve B cells, activated CD4 T cells, cytotoxic CD8 T cells, and NKs.*IRF7* expression decreased through time with decreasing velocity in activated CD4 T cells. For these genes, baseline expression closely resembled day 28 expression from COVID-19 patients except for*IFITM1*, where the regression curve estimated expression to be lower than baseline beyond 15 days following symptom onset.

Among genes whose protein products are related to type I interferon signaling (**Figure 6C**), *HSPB1* (in plasma cells), *CDK6* (in plasma cells), *MNDA* (in XCL+ NKs), and *HLA-DRB5* (in memory B cells) exhibited the minima trend, where expression decreases then increases.*PTPRC* demonstrated the maxima trend, where expression increases then decreases, in cytotoxic CD8 T cells.*CCNE1* (in activated CD4 T cells), *MOV10* (in cytotoxic CD8 T cells), *BCR* (in cytotoxic CD8 T cells), *IGHV3-11* (in activated CD4 T cells), *S1PR3* (in activated CD4 T cells), and *CX3CR1* (in cytotoxic CD8 T cells) showed decreasing expression with decreasing expression velocity through time.*LY6E* (in cytotoxic CD8 T cells, NKs, and naïve T cells), *ICAM2* (in activated CD4 T cells), and *LTB* (in activated CD4 T cells) demonstrated a linear decrease in expression. *NR4A2* (in cytotoxic CD8 T cells), *BLNK* (in NKs), and *TNFRSF13C* (in activated CD4 T cells) showed increasing expression with increasing expression velocity through time. For this set of genes, *CDK6, NR4A1, ICAM2, MNDA, BLNK, CX3CR1*, and *TNFRSF13C* expression did not appear to return to baseline after 28 days.

Prior to metacell analysis by cell type, we also performed the same regression-based time series analysis on all sMetacells (irrespective of cell type) together. With the same R^2^ cutoff of 0.5 or higher and FDR corrected p-value < 0.05, we yielded one significant gene,*ISG15* (**Figure 6D**). The ANOVA p-value for this gene was 8.7e-62 while the R^2^ was 0.55.

## Discussion

### SEACells algorithm generates metacells providing statistical robustness for low replicate time series analysis

In this study, we demonstrate that metacells from the SEACells algorithm (sMetacells) can be used as replicates for time series analysis. Applying it to a COVID-19 scRNA-seq data, we were able to obtain metacells that retained cell-type heterogeneity through time that appear to capture biological variances among individual patients. Despite a similar number of replicates and total cells assigned to metacells, metareplicates from the SEACells algorithm seem less prone to overfitting than those from the rMetacell method, suggesting that the retention of cell type heterogeneity could be important for decreasing overfitting when performing regression on scRNA-seq time series data. sMetacells also maintained a high degree of cell-type purity, enabling us to study expression trends for individual PBMC cell types. As such, our result suggests that this method provides a way to increase statistical power when performing quadratic regression that would otherwise be impossible due to too few replicates. In the absence of this method, pseudobulking led to overfitting, a problem thoroughly defined by Xue Ying [33], which yielded a low number of DEGs with little biological insight. We did not systematically compare the metacells from other algorithms because the SEACells paper has already demonstrated its outperformance to other software [9]. With sMetacells, we were able to obtain a list of significant DEGs for PBMC cell types through time with biological relevance to SARS-CoV-2 infection. Activated CD4 T cells contained the greatest number of significant genes, further validating the reliability of using the SEACells algorithm for time series analysis given CD4 T cells’ critical involvement in response to SARS-CoV-2 infection [34–37].

### ISG15 expression changes significantly through time in the PBMCs

When all PBMC sMetacells were analyzed without using cell type information, we found that*ISG15* was the only gene showing a significant decrease in expression through time. It also exhibited decreasing expression velocity through the 28^th^ day after symptom onset. ISG15 is one of many ISGs that respond to IFN-I to establish an antiviral response [38] and exacerbates inflammation following release from macrophages infected with SARS-CoV-2 [39, 40]. The combination of these findings and this gene’s significance in our analysis further establishes ISG15 as an important part of the immunesystem’s response to SARS-CoV-2. We show that, following infection,*ISG15* expression is initially high 3 days after symptom onset then decreases through day 28 of symptoms. Gene expression velocity also decreases, as is evidenced by the decreasing slope of the line tangent to the fitted expression curve (its derivative) through time. This makes sense since a higher degree of inflammation occurs early in infection when viral load is high then decreases as SARS-CoV-2 is cleared [41].

In the SEACells paper, the authors found that*ISG15* expression was upregulated in CD4 T cells through approximately 10 days after symptom onset and increased again at approximately day 13. Conversely, we found that *ISG15* expression in CD4 T cells decreased continuously with decreasing velocity through approximately 25 days before returning to baseline. This difference could be due to patient cohorts or technical reasons. The SEACells authors constructed metacells from cells of all time points and then determined pseudotime of a metacell based on relative abundance of cells comprising certain time points, and their day 13 metacell was enriched in severe COVID-19 patient cells [9]. We constructed metacells using cells in each of the 10 time points separately. The difference between our results and theirs in relation to *ISG15* may be attributable to continued*ISG15* expression in severe COVID-19 patients. Nevertheless, because of its association with inflammation and disease severity, it will be interesting to study in the future whether changes in expression velocity of*ISG15* would lead to differences in disease severity. This could also be taken a step further to determine whether*ISG15* expression differs between those with and without long COVID-19 symptoms.

### Metacell time series analysis implies that PBMCs and type II pneumocytes share similar SARS-CoV-2 response pathways

Among 165 genes with significant changes in expression through time, the protein products of 89 formed three main clusters within an interaction network generated with STRING. Within these three clusters, 15 genes related to translation, seven related to type I interferon signaling, and one related to cell surface receptor signaling were already annotated in the KEGG COVID-19 pathway. Although this pathway outlines type II pneumocyte response to SARS-CoV-2 and downstream effector cell activation, its significant overlap with our DEGs suggests that despite being non-susceptible to SARS-CoV2 infection [10, 29], PBMCs may undergo a similar response to the virus as type II pneumocytes. PBMCs have been found to induce transcription of interferon-stimulated genes, such as*ISG15* mentioned above, via JAK/STAT signaling upon exposure to SARS-CoV-2 [29]. The KEGG COVID-19 pathway has multiple JAK/STAT signaling cascades that are induced by various cytokines [28]. It may be the case that these same pathways are activated in PBMC response to global cytokine release upon initial infection with SARS-CoV-2.

### Metacell time series analysis implicates new genes not well described in COVID-19 literature

Among the genes not annotated in the KEGG COVID-19 pathway, all have been discussed, albeit most of them only briefly, in previously published COVID-19-related literature.For genes whose protein products are related to translation, EEF2 was previously found to be downregulated in a variety of organ tissue samples from COVID-19 patients compared to controls [42]. We found that EEF2 expression increased through time with decreasing expression velocity in activated CD4 T cells and cytotoxic CD8 T cells.

Earlier time points showed lower expression compared to baseline, suggesting a degree of similarity to the findings from Ghosh et al. Our data suggests that CD4 and CD8 T cells may play an important role in SARS-CoV-2 translation inhibition.

For genes whose protein products are related to type I interferon signaling, IFI27 expression in blood was found to be more highly expressed in patients infected with SARS-CoV-2 as determined via qPCR [43]. Our results show that IFI27 expression decreases significantly through time with decreasing expression velocity before returning to baseline in plasma cells. This suggests that plasma cells may be a large contributor to high IFI27 expression in COVID-19 patient blood. IFIT3 was found to increase in expression through time in SARS-CoV-2 infected mice through 8 days of infection [44]. Interestingly, this conflicts with our results, which show that IFIT3 expression decreases through time with decreasing expression velocity in naïve T cells, naïve B cells, activated CD4 T cells, NKs, and cytotoxic CD8 T cells. IFITM1 was found to inhibit viral RNA production [45] and our data shows a decrease in its expression with decreasing expression velocity in memory B cells. Given IFITM1’s role in inhibiting viral RNA production, a rapid increase in expression of IFITM1 upon exposure to SARS-CoV-2 followed by a gradual decrease through time is expected. We question whether this trend, along with expression velocity, differs depending on previous exposure to SARS-CoV-2 or other coronaviruses. We also notice that IFITM1 expression falls below baseline after 28 days, suggesting potential downregulation of this gene upon clearance of the virus. *IFITM1* has been found to be downregulated following severe influenza infection in mice [46], so we wonder whether our findings could point toward the need to study the differential effects of this gene’s expression in severe and minor COVID-19.

Among genes whose protein products are related to cell surface receptor signaling, LY6E is known to prevent coronavirus fusion [47, 48]. We found that its expression was linear and decreasing in cytotoxic CD8 T cells and NKs but decreasing with decreasing velocity in Naïve T cells. This may point toward high conservation of LY6E’s antiviral activity across different immune cell types. PTPRC (also known as CD45) was found to be more highly expressed in nasopharyngeal cells from SARS-CoV-2 infected patients compared to controls [49]. We found that cytotoxic CD8 T cells exhibit a significant maxima expression trend for this gene, where expression increases then decreases back to baseline by day 28. Since CD45 plays a key role in T cell activation [50], this may suggest that CD8 T cells upregulate this surface protein to mount a strong cytotoxic response over roughly two weeks following COVID-19 symptom onset. ICAM2, a gene whose protein products functions in leukocyte migration [51], was among the 6 most highly up-regulated genes in samples from COVID-19 patient serum [52]. We show that this gene is expressed above baseline and decreases linearly through time; however, its expression continues to decrease below baseline between day 10 and 15 post-symptom onset. This may imply that ICAM2 is down-regulated following viral clearance, perhaps to reestablish a baseline of circulating leukocytes. CX3CR1 expression in NKs has been associated with severe COVID-19 [53]. Our data shows a significant change in expression through time for this gene in cytotoxic CD8 T cells. CX3CR1 expression decreased with decreasing velocity; however, there was also a slight increase in expression after day 20. Additionally, expression did not return to baseline. Given CX3CR1’s association with severe disease and the role of chemokines in inflammation [54], we suggest that this gene may contribute to long COVID-19 symptoms if it continues to be expressed above baseline following virus clearance. Future studies should therefore determine expression trends through time for CX3CR1 in patients with long COVID-19 compared to patients who fully recover.

Although several other significant genes from our analysis have been discussed in literature related to COVID-19, we do not further contribute to their potential role in SARS-CoV-2 infection. We comment only on those where our results are most contributory to previously published materials.

### Limitations

Our study is a proof of concept and generally needs to be applied to more datasets. Furthermore, it needs to be tested more systematically with datasets containing more biological replicates to carefully study the performance difference between true biological replicates and metareplicates. In terms of the relationship of our results to COVID-19, our comparison of day 28 expression to baseline is suboptimal given the low number of metacells per cell type at day 28. We wished to retain expression data through the 28^th^ day after symptom onset, thus we did not perform statistical analysis between day 28 and baseline. Our analysis of expression trends by cell type was also limited by the overall low cell count for certain cell types. This led to low numbers of metacells and subsequent overfitting for these cell types.

## Conclusion

Using the SEACells algorithm to create metacells for time series analysis of COVID-19 data enabled greater statistical power and overcame the limitation of low number of replicates per time point in the original study. We found that*ISG15* expression changed significantly through time when all PBMC cell types were grouped together. This gene demonstrated decreasing expression and decreasing expression velocity through time. For individual cell types, we found many other DEGs through time, which shed new light on our limited knowledge of these genes and their associations with SARS-CoV-2 infection.

## Declarations

### i. Ethics and approval and consent to participate

Not applicable.

### ii. Consent for publication

Not applicable.

### iii. Availability of data and materials

The data analyzed in the current study were described in a previous study (ref 10) and publicly available at the CNGB Nucleotide Sequence Archive (accession number: CNP0001102).

### vi. Authors’ Contributions

K.O. and D.Z. conceived of the experiment. K.O. performed the bioinformatics analysis. K.O. and D.Z. wrote the manuscript.

