## Supplemental Figures and Tables for "Metacell-based differential expression analysis identifies cell type specific temporal gene response programs in COVID-19 patient PBMCs"

### Supplemental Material:

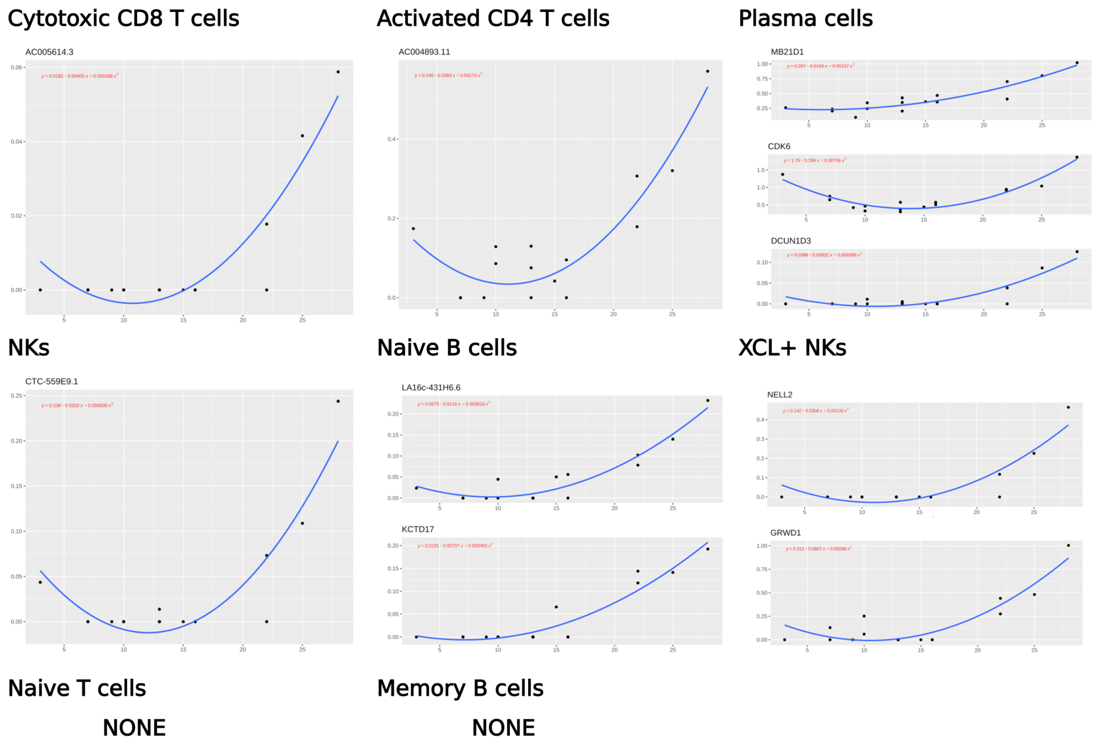

**Supplemental Figure 1: Pseudobulk quadratic regression curves by cell types with formula for genes with FDR adjusted p < 0.05 and $R^2$ > 0.5.** Results for the same 8 cells types as those included in our metacell analysis are shown.

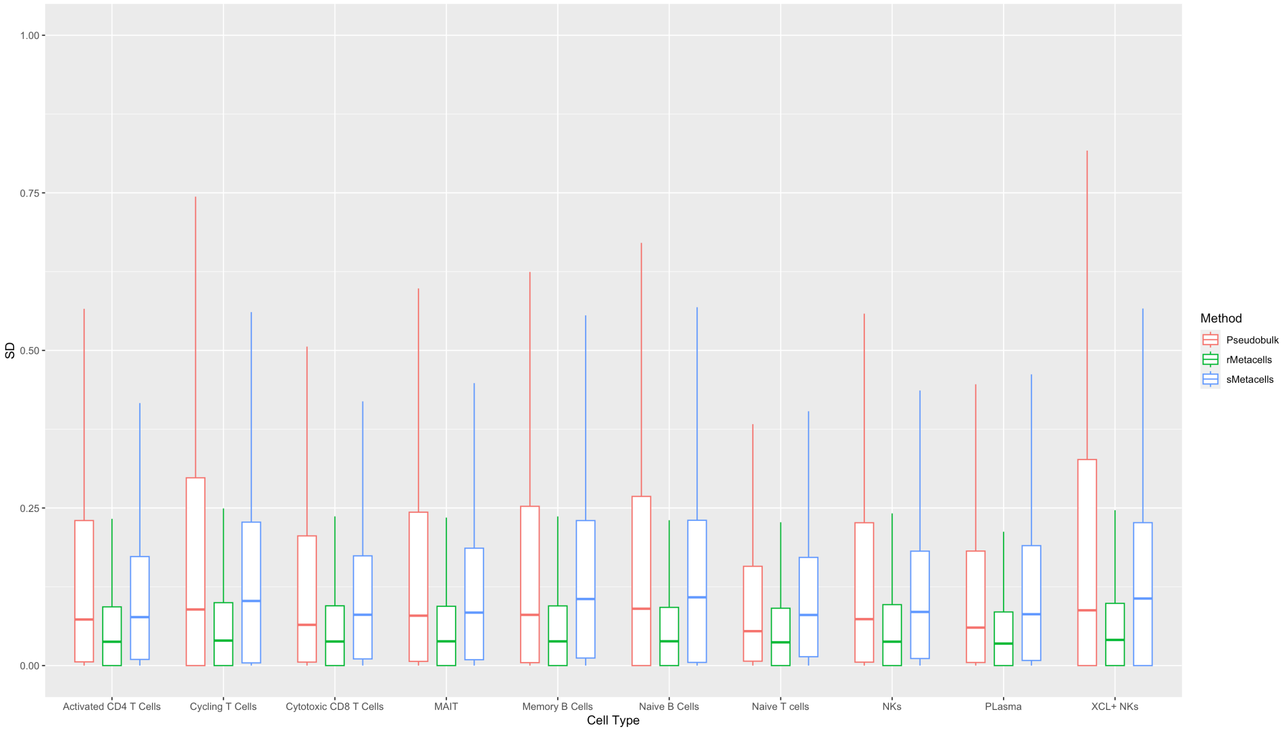

**Supplemental Figure 2: Comparison of SD between rMetacells, sMetacells, and pseudobulked samples.** All SDs were calculated irrespective of time point identification for replicates/metareplicates. Outlier SDs were removed from the plot.

A

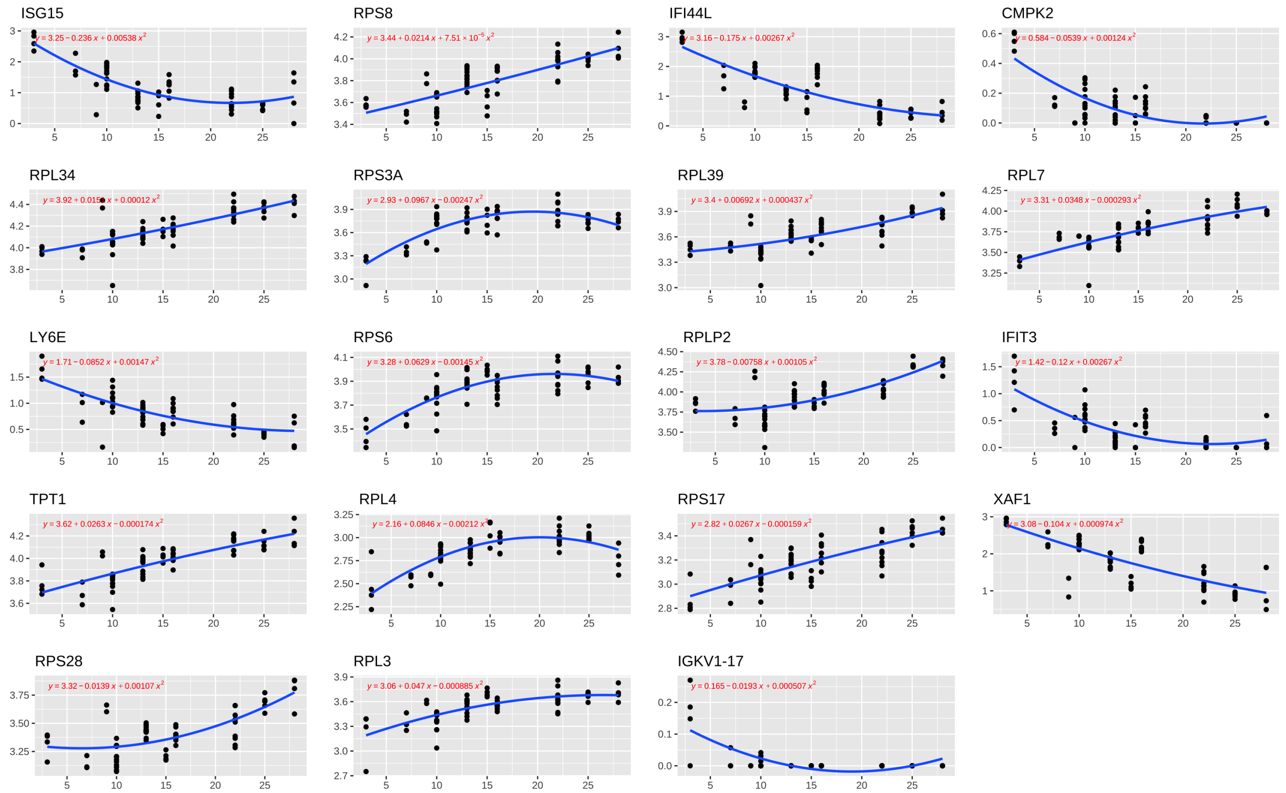
ISG15
RPS8
IFI44L
CMPK2
RPL34
RPS3A
RPL39
RPL7
LY6E
RPS6
RPLP2
IFIT3
TPT1
RPL4
RPS17
XAF1
RPS28
RPL3
IGKV1-17

B

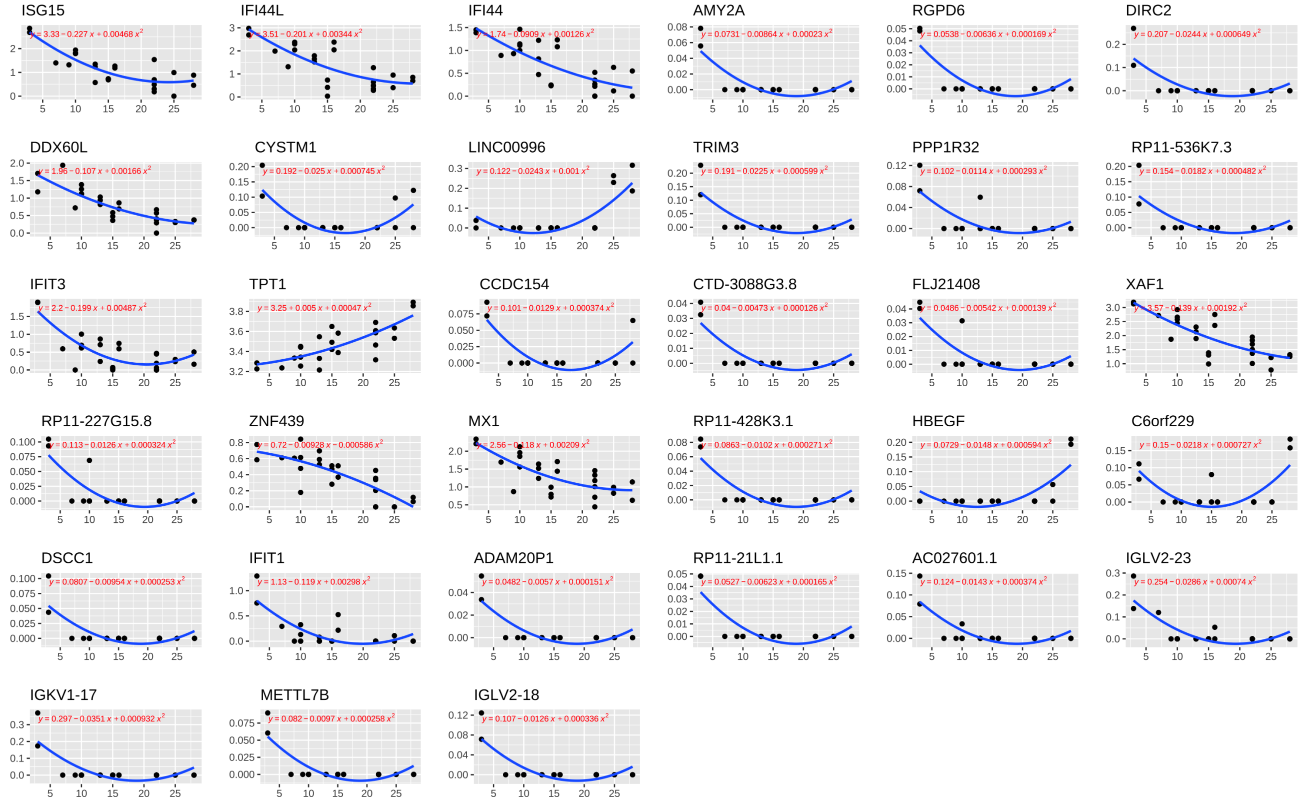
ISG15
IFI44L
IFI44
AMY2A
RGPD6
DIRC2
DDX60L
CYSTM1
LINC00996
TRIM3
PPP1R32
RP11-536K7.3
IFIT3
TPT1
CCDC154
CTD-3088G3.8
FLJ21408
XAF1
RP11-227G15.8
ZNF439
MX1
RP11-428K3.1
HBEGF
C6orf229
DSCC1
IFIT1
ADAM20P1
RP11-21L1.1
AC027601.1
IGLV2-23
IGKV1-17
METTL7B
IGLV2-18

C

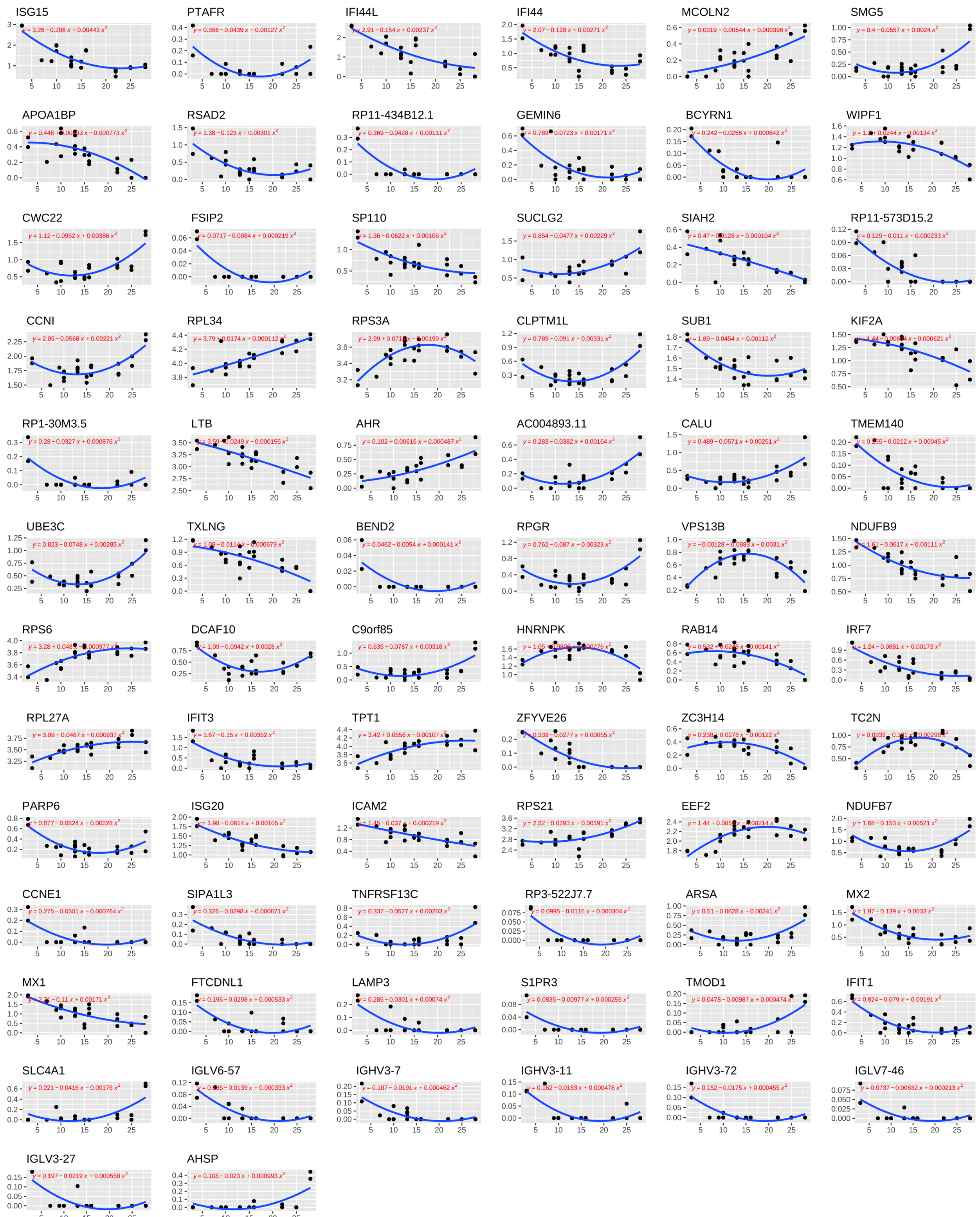
ISG15
PTAFR
IFI44L
IFI44
MCOLN2
SMG5
APOA1BP
RSAD2
RP11-434B12.1
GEMIN6
BCYRN1
WIPF1
CWC22
FSIP2
SP110
SUCLG2
SIAH2
RP11-573D15.2
CCNI
RPL34
RPS3A
CLPTM1L
SUB1
KIF2A
RP1-30M3.5
LTB
AHR
AC004893.11
CALU
TMEM140
UBE3C
TXLNG
BEND2
RPGR
VPS13B
NDUFB9
RPS6
DCAF10
C9orf85
HNRNPK
RAB14
IRF7
RPL27A
IFIT3
TPT1
ZFYVE26
ZC3H14
TC2N
PARP6
ISG20
ICAM2
RPS21
EEF2
NDUFB7
CCNE1
SIPA1L3
TNFRSF13C
RP3-522J7.7
ARSA
MX2
MX1
FTCDNL1
LAMP3
S1PR3
TMOD1
IFIT1
SLC4A1
IGLV6-57
IGHV3-7
IGHV3-11
IGHV3-72
IGLV7-46
IGLV3-27
AHSP

D

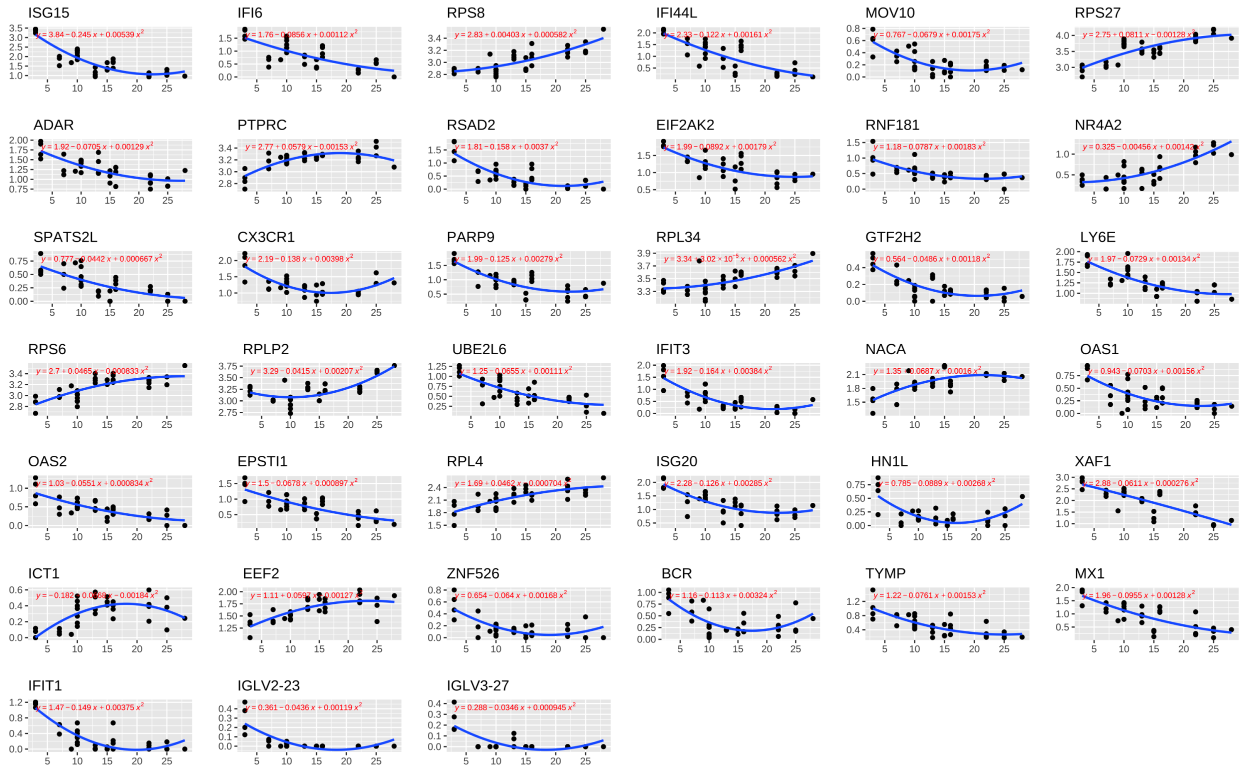
ISG15
IFI6
RPS8
IFI44L
MOV10
RPS27
ADAR
PTPRC
RSAD2
EIF2AK2
RNF181
NR4A2
SPATS2L
CX3CR1
PARP9
RPL34
GTF2H2
LY6E
RPS6
RPLP2
UBE2L6
IFIT3
NACA
OAS1
OAS2
EPSTI1
RPL4
ISG20
HN1L
XAF1
ICT1
EEF2
ZNF526
BCR
TYMP
MX1
IFIT1
IGLV2-23
IGLV3-27

E

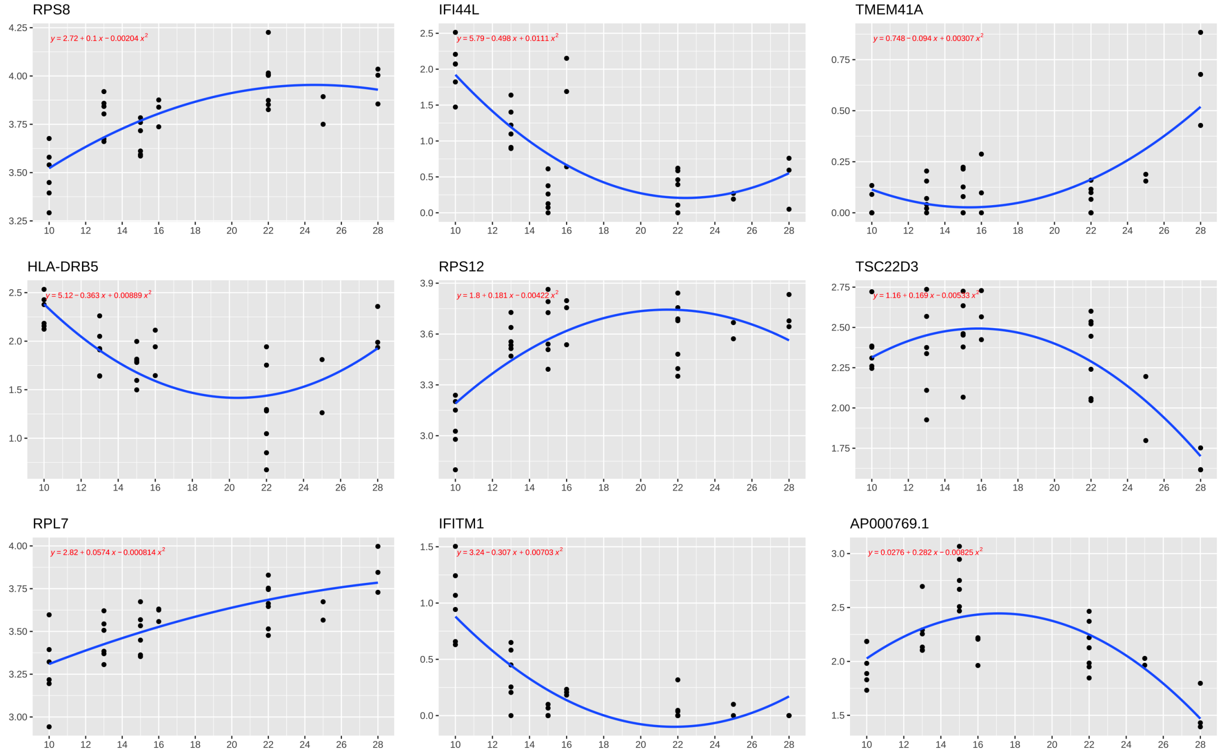
RPS8
IFI44L
TMEM41A
HLA-DRB5
RPS12
TSC22D3
RPL7
IFITM1
AP000769.1

F

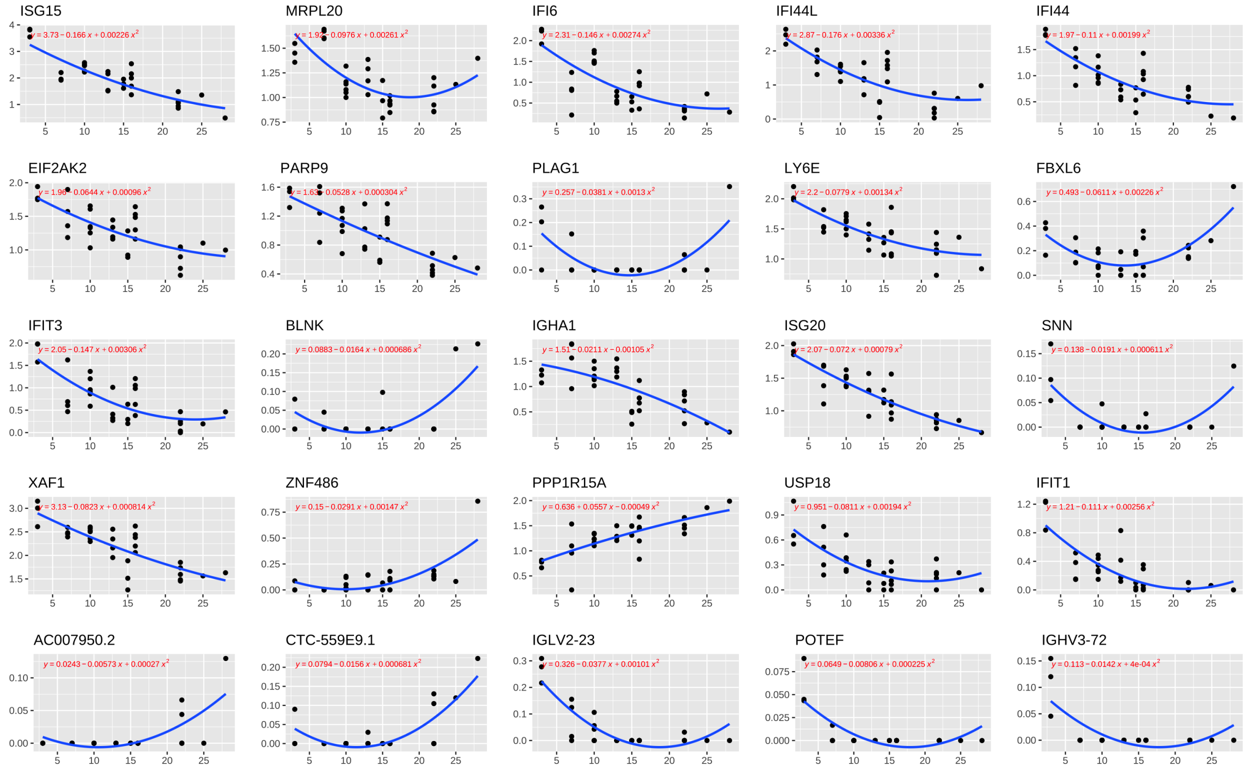
ISG15
MRPL20
IFI6
IFI44L
IFI44
EIF2AK2
PARP9
PLAG1
LY6E
FBXL6
IFIT3
BLNK
IGHA1
ISG20
SNN
XAF1
ZNF486
PPP1R15A
USP18
IFIT1
AC007950.2
CTC-559E9.1
IGLV2-23
POTEF
IGHV3-72

G

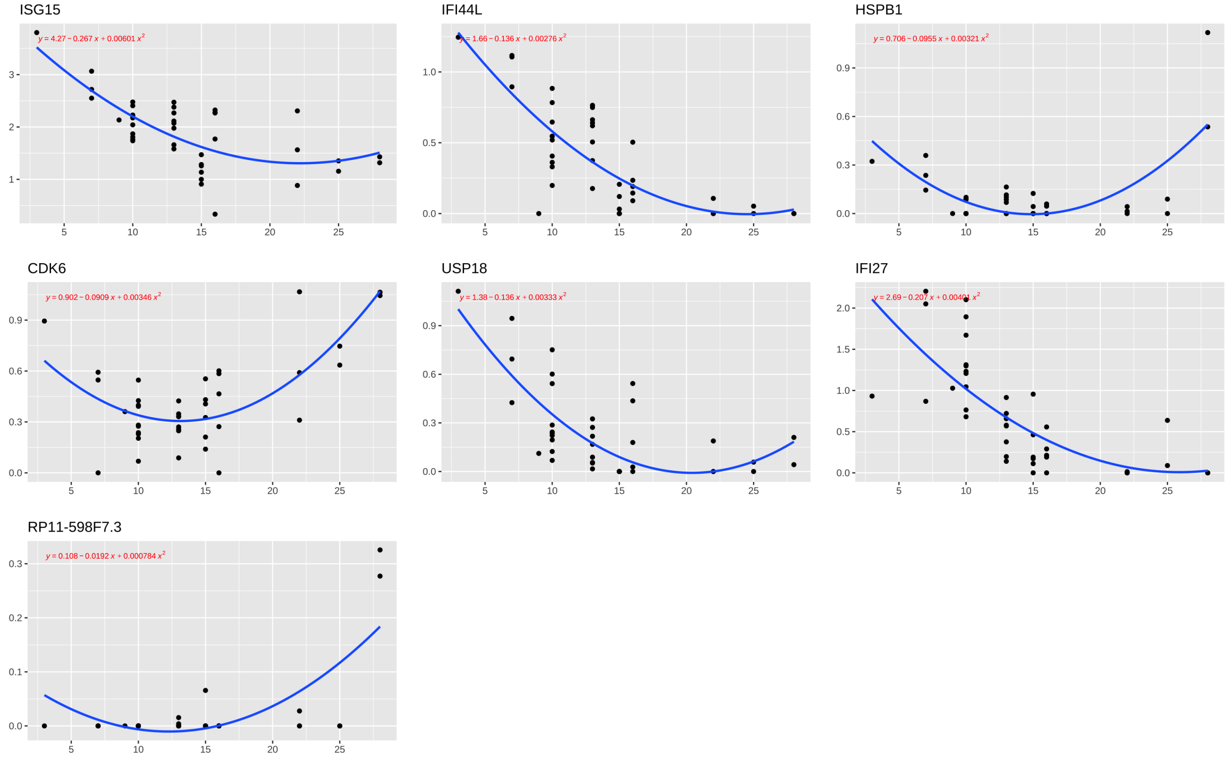
ISG15
IFI44L
HSPB1
CDK6
USP18
IFI27
RP11-598F7.3

H

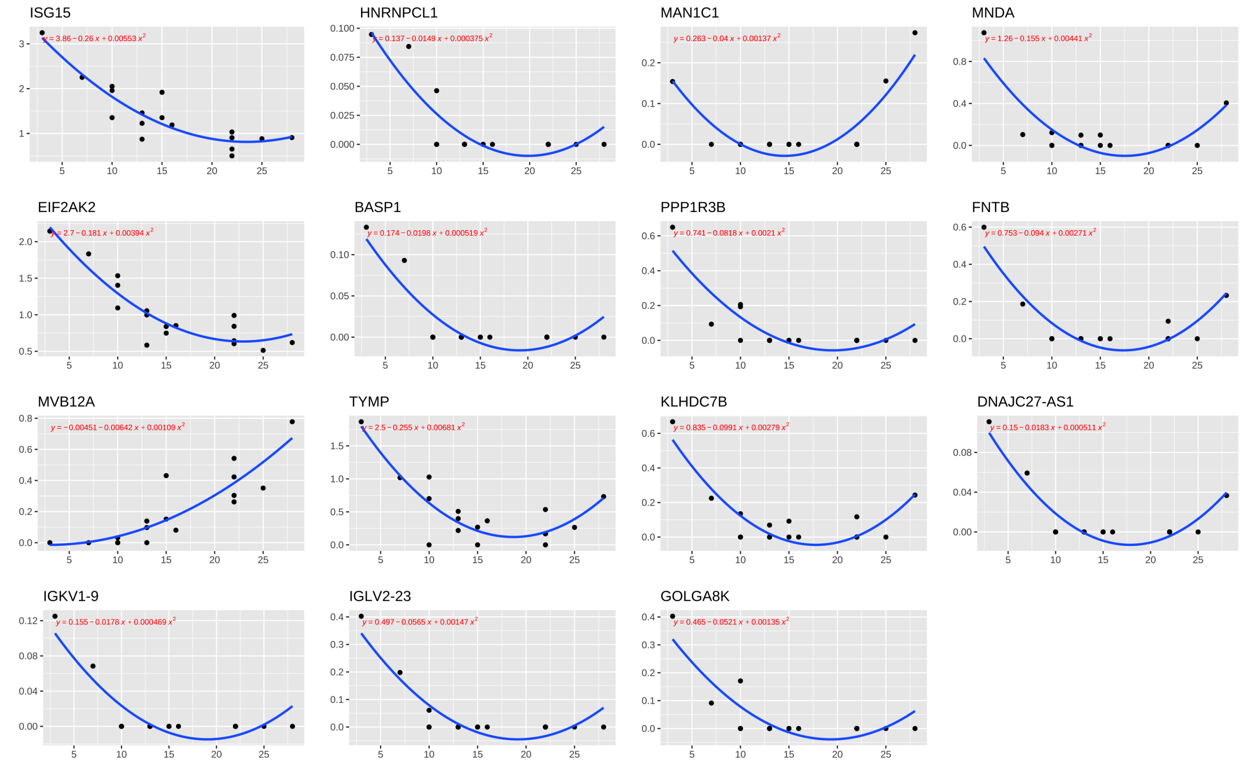

**Supplemental Figure 3: Metacell quadratic regression curves by cell type with formula for genes with FDR adjusted p < 0.05 and $R^2$ > 0.5. A)** Naïve T cells **B)** Naïve B cells **C)** Activated CD4 T cells **D)** Cytotoxic CD8 T cells **E)** Memory B cells **F)** NKs **G)** Plasma cells **H)** XCL+ NKs **A-H)** All x axes represent days since symptom onset while the x axis represents expression.

**A**

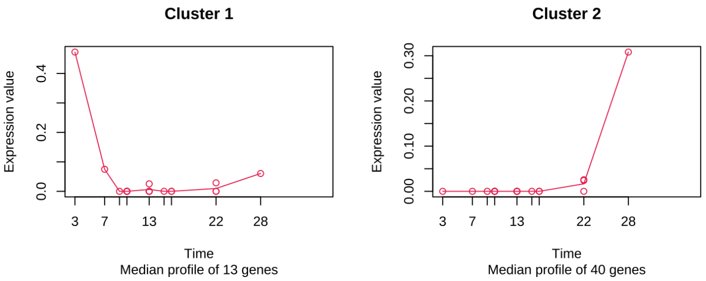

**B**

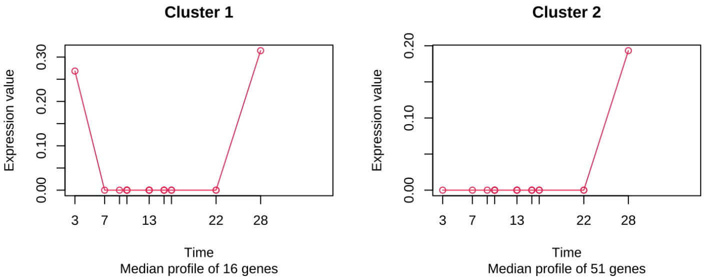

**Supplemental Figure 4: Clustered DEGs from sMetacells for MAITs and cycling T cells using k means clustering A) Clustered expression trends for MAIT cells.** 40 genes in the second cluster are influenced by a single extreme replicate at day 28. **B) Clustered expression trends for cycling T cells.** Both cluster 1 and 2 show extreme outliers influencing expression trends of 67 DEGs.

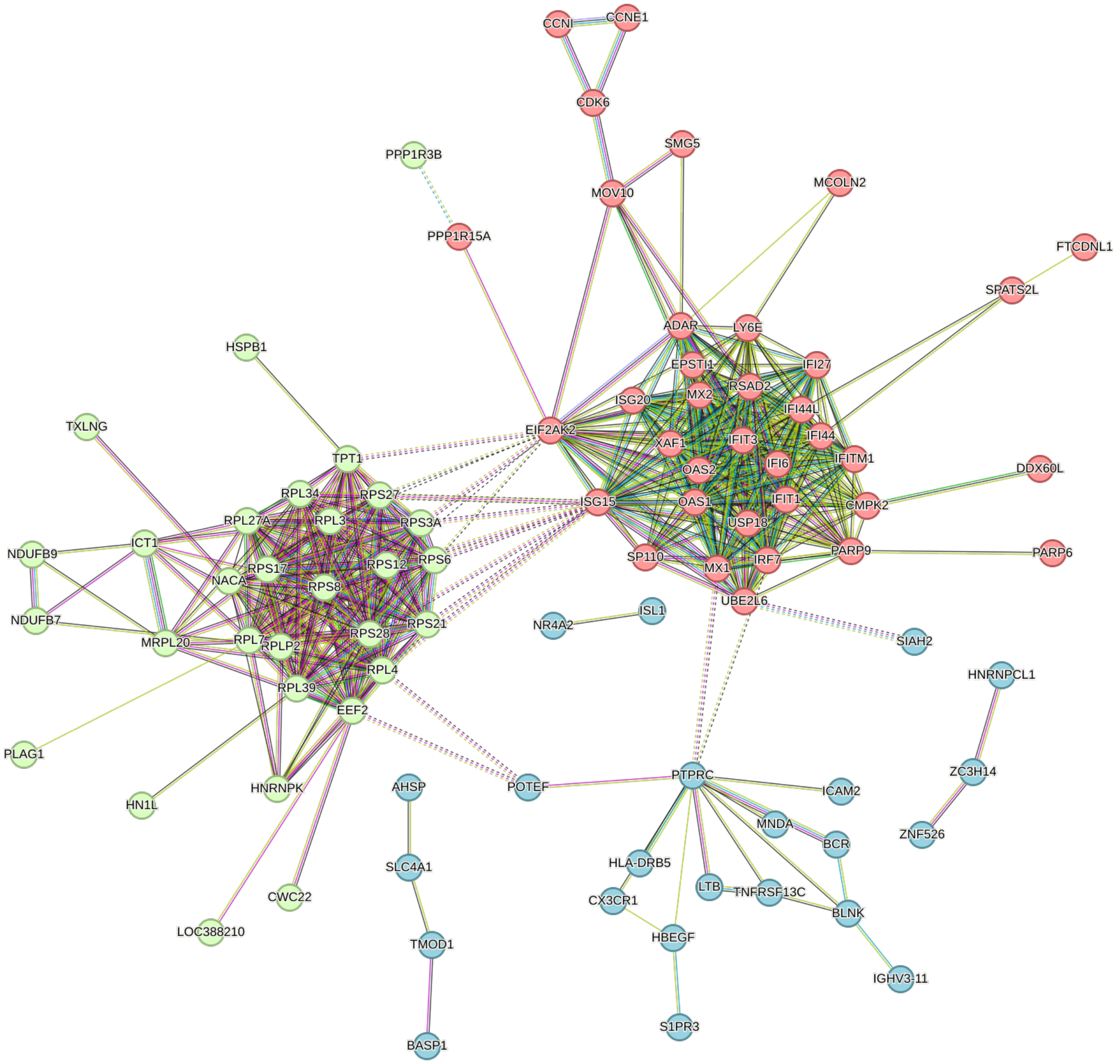

**Supplemental Figure 5:** STRING network colored by K-means clusters.

**Supplemental Table 1: Summary of the average expression standard deviation across cell types.** Only cell types that were compared between s and rMetacells were included. SD was calculated for each gene at each time point for every cell type. The mean SD across all genes was then determined and shown. Values of 0 indicate only one metareplicate for that time point and therefore no variance/SD. “Na” indicates that there were no metareplicates for that time point.

| | **Time Point (Days After Symptom Onset)** | **3** | **7** | **9** | **10** | **13** | **15** | **16** | **22** | **25** | **28** | **Mean** |
|---|---|---|---|---|---|---|---|---|---|---|---|---|
| | All Cell Types | 0.137 | 0.142 | 0.143 | 0.137 | 0.133 | 0.140 | 0.135 | 0.141 | 0.115 | 0.155 | 0.138 |
| | Cytotoxic CD8 T cells | 0.114 | 0.086 | 0.000 | 0.071 | 0.074 | 0.068 | 0.081 | 0.063 | 0.078 | 0.000 | 0.064 |
| | Naïve T cells | 0.107 | 0.093 | 0.081 | 0.076 | 0.073 | 0.082 | 0.074 | 0.086 | 0.073 | 0.119 | 0.086 |
| | NKs | 0.093 | 0.104 | Na | 0.076 | 0.087 | 0.064 | 0.099 | 0.073 | 0.000 | 0.000 | 0.066 |
| | MAIT | 0.000 | 0.000 | 0.000 | 0.099 | 0.084 | 0.000 | 0.078 | 0.069 | Na | 0.000 | 0.037 |
| sMetacells | Activated CD4 T cells | 0.075 | 0.000 | 0.000 | 0.070 | 0.069 | 0.060 | 0.079 | 0.068 | 0.064 | 0.110 | 0.060 |
| | Naïve B | 0.073 | 0.000 | 0.000 | 0.114 | 0.101 | 0.072 | 0.090 | 0.131 | 0.083 | 0.096 | 0.076 |
| | Plasma | 0.000 | 0.115 | 0.000 | 0.094 | 0.090 | 0.093 | 0.105 | 0.096 | 0.073 | 0.091 | 0.076 |
| | Memory B | 0.000 | 0.000 | 0.000 | 0.119 | 0.117 | 0.093 | 0.100 | 0.118 | 0.062 | 0.120 | 0.073 |
| | XCL+ NKs | 0.000 | 0.000 | Na | 0.105 | 0.112 | 0.081 | 0.000 | 0.102 | 0.000 | 0.000 | 0.044 |
| | Cycling T cells | 0.000 | 0.000 | 0.000 | 0.125 | 0.113 | 0.096 | 0.105 | 0.095 | Na | 0.000 | 0.059 |
| | All Cell Types | 0.049 | 0.049 | 0.044 | 0.049 | 0.050 | 0.051 | 0.053 | 0.049 | 0.048 | 0.049 | 0.049 |
| | Cytotoxic CD8 T cells | 0.043 | 0.043 | 0.026 | 0.047 | 0.047 | 0.047 | 0.051 | 0.048 | 0.040 | 0.047 | 0.044 |
| | Naïve T cells | 0.043 | 0.042 | 0.035 | 0.045 | 0.047 | 0.043 | 0.049 | 0.044 | 0.044 | 0.046 | 0.044 |
| | NKs | 0.044 | 0.044 | 0.029 | 0.046 | 0.048 | 0.047 | 0.051 | 0.049 | 0.042 | 0.048 | 0.045 |
| | MAIT | 0.039 | 0.039 | 0.032 | 0.046 | 0.047 | 0.041 | 0.043 | 0.047 | 0.037 | 0.044 | 0.042 |
| rMetacells | Activated CD4 T cells | 0.040 | 0.036 | 0.031 | 0.045 | 0.047 | 0.044 | 0.041 | 0.046 | 0.043 | 0.046 | 0.042 |
| | Naïve B | 0.042 | 0.034 | 0.030 | 0.044 | 0.046 | 0.045 | 0.018 | 0.042 | 0.036 | 0.042 | 0.038 |
| | Plasma | 0.030 | 0.030 | 0.030 | 0.039 | 0.044 | 0.039 | 0.040 | 0.042 | 0.041 | 0.041 | 0.038 |
| | Memory B | 0.041 | 0.006 | 0.024 | 0.044 | 0.045 | 0.047 | 0.032 | 0.046 | 0.045 | 0.044 | 0.037 |
| | XCL+ NKs | 0.037 | 0.039 | Na | 0.044 | 0.047 | 0.045 | 0.045 | 0.051 | 0.042 | 0.042 | 0.043 |
| | Cycling T cells | 0.014 | 0.033 | Na | 0.045 | 0.049 | 0.048 | 0.035 | 0.043 | 0.000 | 0.034 | 0.033 |

**Supplemental Table 2: Summary of the numbers of patient samples, single cells, and SEACells-generated metacells.** A, the number of samples and cells for each of the 10 time points. B, the number of metacells representing each cell type per time point.

| | **Time Point (Days After Symptom Onset)** | **3** | **7** | **9** | **10** | **13** | **15** | **16** | **22** | **25** | **28** | **Total** |
|---|---|---|---|---|---|---|---|---|---|---|---|---|
| A | # Samples/patients | 1 | 2 | 1 | 2 | 3 | 1 | 2 | 2 | 1 | 1 | 16 |
| | # Cells | 1240 | 1512 | 528 | 5433 | 4627 | 2877 | 3267 | 4272 | 1488 | 531 | 25775 |
| B | Naïve T cells | 4 | 3 | 2 | 12 | 12 | 4 | 8 | 9 | 6 | 4 | 64 |
| | Activated CD4 T cells | 2 | 1 | 1 | 3 | 5 | 2 | 3 | 3 | 2 | 2 | 24 |
| | NKs | 3 | 4 | 0 | 6 | 4 | 3 | 5 | 5 | 1 | 1 | 32 |
| | Naïve B cells | 2 | 1 | 1 | 4 | 3 | 3 | 2 | 6 | 2 | 2 | 26 |
| | MAIT cells | 1 | 1 | 1 | 5 | 4 | 1 | 2 | 3 | 0 | 1 | 19 |
| | Cytotoxic CD8 T cells | 4 | 3 | 1 | 7 | 4 | 3 | 4 | 5 | 3 | 1 | 35 |
| | Memory B cells | 1 | 1 | 1 | 6 | 6 | 6 | 3 | 7 | 2 | 3 | 36 |
| | Cycling T cells | 1 | 1 | 1 | 6 | 5 | 4 | 3 | 2 | 0 | 1 | 24 |
| | XCL+ NKs | 1 | 1 | 0 | 3 | 3 | 2 | 1 | 4 | 1 | 1 | 17 |
| | Plasma cells | 1 | 3 | 1 | 10 | 8 | 6 | 5 | 3 | 2 | 2 | 41 |
| | Monocytes | 0 | 1 | 1 | 1 | 1 | 1 | 1 | 7 | 0 | 1 | 14 |
| | Stem cells | 0 | 0 | 0 | 2 | 1 | 1 | 1 | 1 | 0 | 0 | 6 |
| | Megakaryocytes | 0 | 0 | 0 | 1 | 0 | 2 | 0 | 0 | 0 | 0 | 3 |
| | Cycling Plasma | 0 | 0 | 0 | 4 | 2 | 1 | 1 | 2 | 1 | 0 | 11 |
| | DCs | 0 | 0 | 0 | 0 | 2 | 1 | 1 | 3 | 0 | 1 | 8 |
| | Total # Metacells | 20 | 20 | 10 | 70 | 60 | 40 | 40 | 60 | 20 | 10 | 360 |

**Supplemental Table 3: All genes deemed significant using maSigPro by cell type.** FDR adjusted p-values are all less than 0.05 while $R^2$ is greater than 0.5. For each gene and cell type, the overall expression trend pattern is also shown.

| Pattern | Cell Type | Gene | -log(p value) | R2 | ANOVA p value |
|---|---|---|---|---|---|
| Increasing, constant | Naive T | RPL7 | 16.37863664 | 0.6828868 | 4.18E-17 |
| Increasing, constant | Naive T | TPT1 | 16.00242853 | 0.6739667 | 9.94E-17 |
| Increasing, constant | Naive T | RPS8 | 14.76655535 | 0.6428808 | 1.71E-15 |
| Decreasing, ↓ velocity | Naive T | IFI44L | 14.69832664 | 0.6703238 | 2.00E-15 |
| Increasing, constant | Naive T | RPS17 | 14.25199379 | 0.6290911 | 5.60E-15 |
| Increasing, ↓ velocity | Naive T | RPS6 | 13.50811187 | 0.6393291 | 3.10E-14 |
| Increasing, constant | Naive T | RPL34 | 13.45347831 | 0.6066459 | 3.52E-14 |
| Decreasing, constant | Naive T | XAF1 | 13.41885725 | 0.6056431 | 3.81E-14 |
| Increasing, ↓ velocity | Naive T | RPS3A | 13.28022097 | 0.6330703 | 5.25E-14 |
| Decreasing, ↓ velocity | Naive T | ISG15 | 12.98834245 | 0.6248951 | 1.03E-13 |
| Increasing, ↑ velocity | Naive T | RPLP2 | 12.61481518 | 0.5816333 | 2.43E-13 |
| Decreasing, ↓ velocity | Naive T | CMPK2 | 12.24918647 | 0.6033684 | 5.63E-13 |
| Increasing, ↓ velocity | Naive T | RPL4 | 12.03633686 | 0.5969434 | 9.20E-13 |
| Increasing, ↑ velocity | Naive T | RPL39 | 11.79606789 | 0.5557109 | 1.60E-12 |
| Decreasing, ↓ velocity | Naive T | LY6E | 11.64865484 | 0.5849725 | 2.25E-12 |
| Increasing, ↑ velocity | Naive T | RPS28 | 11.52853857 | 0.5469046 | 2.96E-12 |
| Decreasing, ↓ velocity | Cytotoxic CD8 T | ISG15 | 10.88028069 | 0.7910788 | 1.31741E-11 |
| Decreasing, ↓ velocity | Naive T | IFIT3 | 10.55630403 | 0.5492957 | 2.78E-11 |
| Decreasing, ↓ velocity | Cytotoxic CD8 T | IFIT1 | 10.07461117 | 0.7653952 | 8.42149E-11 |
| Decreasing, constant | NK | ISG20 | 9.988230004 | 0.7565666 | 1.02747E-10 |
| Increasing, ↓ velocity | Naive T | RPL3 | 9.551191841 | 0.5137651 | 2.81E-10 |
| Decreasing, ↓ velocity | Naive T | IGKV1-17 | 9.304003867 | 0.5046061 | 4.97E-10 |
| Decreasing, constant | Cytotoxic CD8 T | XAF1 | 9.175062202 | 0.6898753 | 6.68248E-10 |
| Increasing, ↑ velocity | Cytotoxic CD8 T | NR4A2 | 8.954302359 | 0.6802974 | 1.11096E-09 |
| Decreasing, ↓ velocity | NK | IGLV2-23 | 8.799737346 | 0.7527584 | 1.58585E-09 |
| Decreasing, constant | NK | ISG15 | 8.772547035 | 0.7072286 | 1.68831E-09 |
| Increasing, ↑ velocity | Cytotoxic CD8 T | RPS8 | 8.286920593 | 0.6495437 | 5.16511E-09 |
| Decreasing, constant | Cytotoxic CD8 T | IFI44L | 8.107769841 | 0.6408069 | 7.80244E-09 |
| Increasing, constant | Cytotoxic CD8 T | RPS27 | 7.89583429 | 0.6301963 | 1.27106E-08 |
| Decreasing, ↓ velocity | Plasma | IFI44L | 7.864587454 | 0.6144557 | 1.36588E-08 |
| Decreasing, ↓ velocity | Cytotoxic CD8 T | RSAD2 | 7.73048799 | 0.6712661 | 1.86E-08 |
| Decreasing, ↓ velocity | Cytotoxic CD8 T | IFIT3 | 7.688437483 | 0.6692707 | 2.0491E-08 |
| Decreasing, constant | Cytotoxic CD8 T | EPSTI1 | 7.600860738 | 0.6149183 | 2.50691E-08 |
| Decreasing, ↓ velocity | Mem B | IFITM1 | 7.597916844 | 0.6884898 | 2.52396E-08 |
| Decreasing, constant | Cytotoxic CD8 T | MX1 | 7.534865005 | 0.6114168 | 2.91833E-08 |
| Decreasing, constant | NK | XAF1 | 7.471678743 | 0.6435442 | 3.37537E-08 |
| Decreasing, ↓ velocity | Cytotoxic CD8 T | PARP9 | 7.414209332 | 0.6559576 | 3.85293E-08 |

| | | | | | |
|---|---|---|---|---|---|
| Decreasing, ↓ velocity | Cytotoxic CD8 T | IGLV2-23 | 7.279599627 | 0.6492279 | 5.25292E-08 |
| Decreasing, constant | Cytotoxic CD8 T | OAS2 | 7.039302763 | 0.5841183 | 9.13476E-08 |
| Increasing, constant | Cytotoxic CD8 T | RPS6 | 6.968725279 | 0.5800823 | 1.07467E-07 |
| Increasing, ↑ velocity | Plasma | RP11-598F7.3 | 6.902763867 | 0.5667918 | 1.25094E-07 |
| Increasing, ↑ velocity | NK | CTC-559E9.1 | 6.899144075 | 0.6656534 | 1.26141E-07 |
| Decreasing, constant | Cytotoxic CD8 T | LY6E | 6.891453918 | 0.57562 | 1.28394E-07 |
| Decreasing, ↓ velocity | Plasma | USP18 | 6.735017931 | 0.5578951 | 1.8407E-07 |
| Decreasing, constant | Naïve B | DDX60L | 6.713932863 | 0.6835548 | 1.93227E-07 |
| Decreasing, constant | Cytotoxic CD8 T | UBE2L6 | 6.690856984 | 0.5638212 | 2.03771E-07 |
| Decreasing, constant | NK | IFI44 | 6.66740078 | 0.5976059 | 2.1508E-07 |
| Decreasing, constant | Cytotoxic CD8 T | ADAR | 6.583301424 | 0.5573652 | 2.61035E-07 |
| Min Parabolic | Plasma | CDK6 | 6.569326186 | 0.5489279 | 2.69571E-07 |
| Decreasing, ↓ velocity | Naïve B | ISG15 | 6.529124589 | 0.7294484 | 2.95716E-07 |
| Min Parabolic | Plasma | HSPB1 | 6.528624373 | 0.5466974 | 2.96057E-07 |
| Decreasing, constant | NK | PARP9 | 6.445133162 | 0.5839298 | 3.58812E-07 |
| Increasing, constant | Cytotoxic CD8 T | RPL4 | 6.393656154 | 0.545757 | 4.03965E-07 |
| Decreasing, ↓ velocity | XCL+ NKs | DNAJC27-AS1 | 6.311889367 | 0.8745989 | 4.87653E-07 |
| Decreasing, constant | Cytotoxic CD8 T | IFI6 | 6.295010801 | 0.5396038 | 5.06978E-07 |
| Max Parabolic | Cytotoxic CD8 T | PTPRC | 6.279631706 | 0.5949366 | 5.25253E-07 |
| Decreasing, ↓ velocity | NK | IFIT1 | 6.276158642 | 0.6308852 | 5.2947E-07 |
| Decreasing, constant | NK | LY6E | 6.262527542 | 0.5723593 | 5.46352E-07 |
| Decreasing, ↓ velocity | Cytotoxic CD8 T | CX3CR1 | 6.24659828 | 0.5930064 | 5.66763E-07 |
| Decreasing, ↓ velocity | Plasma | IFI27 | 6.246165552 | 0.5309119 | 5.67328E-07 |
| Decreasing, ↓ velocity | Cytotoxic CD8 T | ISG20 | 6.242009577 | 0.5927375 | 5.72783E-07 |
| Decreasing, ↓ velocity | Naïve B | RP11-21L1.1 | 6.239549846 | 0.7132982 | 5.76037E-07 |
| Decreasing, ↓ velocity | Naïve B | RGPD6 | 6.23329113 | 0.7129387 | 5.84398E-07 |
| Increasing, constant | Mem B | RPL7 | 6.233264675 | 0.5584398 | 5.84434E-07 |
| Decreasing, ↓ velocity | Cytotoxic CD8 T | TYMP | 6.19780047 | 0.5901382 | 6.34161E-07 |
| Decreasing, ↓ velocity | Cytotoxic CD8 T | GTF2H2 | 6.187644101 | 0.5895387 | 6.49166E-07 |
| Decreasing, ↓ velocity | Cytotoxic CD8 T | MOV10 | 6.179742894 | 0.5890717 | 6.61085E-07 |
| Decreasing, ↓ velocity | Naïve B | RP11-428K3.1 | 6.176866754 | 0.7096772 | 6.65477E-07 |
| Increasing, constant | NK | PPP1R15A | 6.131735956 | 0.5638821 | 7.38353E-07 |
| Decreasing, ↓ velocity | Mem B | IFI44L | 6.076932274 | 0.6065666 | 8.3766E-07 |
| Decreasing, ↓ velocity | Activated CD4 T | RP3-522J7.7 | 6.069746906 | 0.7358024 | 8.51634E-07 |
| Decreasing, ↓ velocity | Naïve B | CTD-3088G3.8 | 6.068836166 | 0.703329 | 8.53422E-07 |
| Decreasing, ↑ velocity | Naïve B | ZNF439 | 6.038734046 | 0.6406502 | 9.14673E-07 |
| Decreasing, constant | Cytotoxic CD8 T | SPATS2L | 6.037796789 | 0.5231816 | 9.16649E-07 |
| Decreasing, ↓ velocity | Cytotoxic CD8 T | EIF2AK2 | 6.000985837 | 0.5783633 | 9.97733E-07 |
| Decreasing, ↓ velocity | NK | POTEF | 5.996766584 | 0.6141399 | 1.00747E-06 |
| Decreasing, ↓ velocity | Plasma | ISG15 | 5.985826111 | 0.5158761 | 1.03318E-06 |

| | | | | | |
|---|---|---|---|---|---|
| Increasing, ↓ velocity | Cytotoxic CD8 T | EEF2 | 5.980829686 | 0.5771385 | 1.04513E-06 |
| Decreasing, ↓ velocity | Activated CD4 T | ISG20 | 5.978318779 | 0.7304519 | 1.05119E-06 |
| Increasing, constant | Activated CD4 T | RPL34 | 5.966809539 | 0.6682814 | 1.07942E-06 |
| Decreasing, ↓ velocity | Activated CD4 T | FSIP2 | 5.956123947 | 0.7291367 | 1.10631E-06 |
| Decreasing, constant | NK | IFI44L | 5.923523403 | 0.5500528 | 1.19255E-06 |
| Increasing, ↓ velocity | Mem B | RPS8 | 5.887321617 | 0.594947 | 1.29622E-06 |
| Decreasing, ↓ velocity | Naïve B | AMY2A | 5.885264616 | 0.6922218 | 1.30237E-06 |
| Increasing, ↑ velocity | Naïve B | TPT1 | 5.861515726 | 0.6284844 | 1.37558E-06 |
| Decreasing, ↓ velocity | Activated CD4 T | IFIT1 | 5.787539648 | 0.7189357 | 1.63102E-06 |
| Decreasing, constant | Naïve B | XAF1 | 5.785779176 | 0.6231646 | 1.63765E-06 |
| Decreasing, ↓ velocity | Naïve B | METTL7B | 5.770514492 | 0.6850685 | 1.69623E-06 |
| Decreasing, ↑ velocity | NK | IGHA1 | 5.759853035 | 0.5388878 | 1.73839E-06 |
| Min Parabolic | Naïve B | CCDC154 | 5.745250894 | 0.6834714 | 1.79783E-06 |
| Decreasing, ↓ velocity | Activated CD4 T | RP11-434B12.1 | 5.730756355 | 0.7154139 | 1.85885E-06 |
| Increasing, ↑ velocity | Cytotoxic CD8 T | RPL34 | 5.721980978 | 0.5022521 | 1.89679E-06 |
| Decreasing, ↓ velocity | Activated CD4 T | ZFYVE26 | 5.676833529 | 0.7120287 | 2.10459E-06 |
| Decreasing, ↓ velocity | Activated CD4 T | IGLV6-57 | 5.671790682 | 0.7117101 | 2.12917E-06 |
| Decreasing, ↓ velocity | Naïve B | IGLV2-23 | 5.613714399 | 0.6750243 | 2.4338E-06 |
| Min Parabolic | XCL+ NKs | FNTB | 5.6010971 | 0.8415679 | 2.50555E-06 |
| Decreasing, ↓ velocity | XCL+ NKs | IGLV2-23 | 5.577408191 | 0.8403285 | 2.64601E-06 |
| Decreasing, ↓ velocity | Naïve B | ADAM20P1 | 5.576514756 | 0.6725947 | 2.65146E-06 |
| Decreasing, ↓ velocity | Cytotoxic CD8 T | IGLV3-27 | 5.549968401 | 0.5500886 | 2.81859E-06 |
| Increasing, ↓ velocity | Cytotoxic CD8 T | NACA | 5.539232315 | 0.549393 | 2.88913E-06 |
| Increasing, ↑ velocity | Activated CD4 T | UBE3C | 5.517058849 | 0.70176 | 3.04047E-06 |
| Increasing, ↑ velocity | Naïve B | LINC00996 | 5.475830367 | 0.6659274 | 3.34326E-06 |
| Decreasing, ↓ velocity | Activated CD4 T | IGHV3-72 | 5.474236027 | 0.6989461 | 3.35555E-06 |
| Decreasing, ↓ velocity | XCL+ NKs | EIF2AK2 | 5.472447428 | 0.8347194 | 3.3694E-06 |
| Increasing, ↑ velocity | XCL+ NKs | MVB12A | 5.464195409 | 0.7722946 | 3.43403E-06 |
| Decreasing, constant | NK | EIF2AK2 | 5.458491023 | 0.5176345 | 3.47944E-06 |
| Increasing, ↑ velocity | Cytotoxic CD8 T | RPLP2 | 5.435961335 | 0.5426461 | 3.6647E-06 |
| Decreasing, ↓ velocity | Naïve B | AC027601.1 | 5.417915605 | 0.662031 | 3.82019E-06 |
| Decreasing, ↓ velocity | Naïve B | IGLV2-18 | 5.402612658 | 0.6609938 | 3.95719E-06 |
| Min Parabolic | NK | SNN | 5.39782168 | 0.5756381 | 4.00109E-06 |
| Decreasing, ↓ velocity | Activated CD4 T | MX2 | 5.342798769 | 0.6901425 | 4.54152E-06 |
| Decreasing, constant | Activated CD4 T | LTB | 5.305236203 | 0.6201512 | 4.95181E-06 |
| Decreasing, ↓ velocity | XCL+ NKs | ISG15 | 5.294759023 | 0.8247711 | 5.07272E-06 |
| Min Parabolic | Cytotoxic CD8 T | HN1L | 5.264689794 | 0.5312332 | 5.43639E-06 |
| Decreasing, ↓ velocity | NK | IFIT3 | 5.260532722 | 0.5662848 | 5.48867E-06 |
| Decreasing, constant | NK | IFI6 | 5.253024104 | 0.5026137 | 5.58439E-06 |
| Increasing, ↑ velocity | Activated CD4 T | AC004893.11 | 5.249859947 | 0.6837625 | 5.62523E-06 |

| | | | | | |
|---|---|---|---|---|---|
| Decreasing, ↓ velocity | NK | USP18 | 5.236511509 | 0.5646272 | 5.80081E-06 |
| Decreasing, ↓ velocity | Naïve B | IFIT3 | 5.233414139 | 0.6493123 | 5.84233E-06 |
| Decreasing, ↓ velocity | Naïve B | TRIM3 | 5.176173783 | 0.64527 | 6.6654E-06 |
| Decreasing, ↓ velocity | Naïve B | RP11-227G15.8 | 5.170191195 | 0.6448448 | 6.75785E-06 |
| Increasing, ↑ velocity | Mem B | TMEM41A | 5.134535645 | 0.5453286 | 7.33609E-06 |
| Decreasing, constant | Naïve B | IFI44L | 5.121921917 | 0.5732849 | 7.55228E-06 |
| Min Parabolic | XCL+ NKs | KLHDC7B | 5.103911384 | 0.813418 | 7.87206E-06 |
| Decreasing, constant | Activated CD4 T | NDUFB9 | 5.100530138 | 0.6039396 | 7.93359E-06 |
| Decreasing, ↑ velocity | Activated CD4 T | APOA1BP | 5.092239785 | 0.6032693 | 8.08649E-06 |
| Decreasing, ↓ velocity | Cytotoxic CD8 T | BCR | 5.067617721 | 0.5177482 | 8.5582E-06 |
| Decreasing, ↓ velocity | Cytotoxic CD8 T | ZNF526 | 5.048301545 | 0.5164057 | 8.94743E-06 |
| Increasing, ↓ velocity | Cytotoxic CD8 T | ICT1 | 5.045565092 | 0.5162152 | 9.00399E-06 |
| Increasing, ↑ velocity | Activated CD4 T | AHR | 5.043310537 | 0.5992906 | 9.05085E-06 |
| Decreasing, ↓ velocity | NK | IGHV3-72 | 5.039698645 | 0.5508053 | 9.12644E-06 |
| Decreasing, constant | Naïve B | IFI44 | 5.037133665 | 0.5664758 | 9.1805E-06 |
| Min Parabolic | Mem B | HLA-DRB5 | 5.036365288 | 0.538425 | 9.19676E-06 |
| Decreasing, ↓ velocity | XCL+ NKs | IGKV1-9 | 5.023386773 | 0.8084098 | 9.47574E-06 |
| Decreasing, ↓ velocity | Naïve B | FLJ21408 | 5.014053896 | 0.6335664 | 9.68158E-06 |
| Increasing, ↑ velocity | XCL+ NKs | MAN1C1 | 5.01002583 | 0.8075659 | 9.77179E-06 |
| Decreasing, ↓ velocity | Activated CD4 T | IRF7 | 5.002575081 | 0.6661401 | 9.94088E-06 |
| Decreasing, ↓ velocity | Activated CD4 T | ISG15 | 4.94508029 | 0.661904 | 1.1348E-05 |
| Decreasing, ↑ velocity | Mem B | TSC22D3 | 4.938255451 | 0.5314208 | 1.15278E-05 |
| Decreasing, constant | Activated CD4 T | MX1 | 4.937129334 | 0.5905247 | 1.15577E-05 |
| Decreasing, ↓ velocity | Cytotoxic CD8 T | RNF181 | 4.900625337 | 0.5060182 | 1.25711E-05 |
| Decreasing, ↓ velocity | Naïve B | IGKV1-17 | 4.900248248 | 0.6251207 | 1.25821E-05 |
| Min Parabolic | NK | PLAG1 | 4.893052383 | 0.5402221 | 1.27923E-05 |
| Decreasing, ↓ velocity | Cytotoxic CD8 T | OAS1 | 4.88939009 | 0.5052189 | 1.29006E-05 |
| Decreasing, ↓ velocity | NK | MRPL20 | 4.86992077 | 0.5385301 | 1.34921E-05 |
| Increasing, ↑ velocity | NK | ZNF486 | 4.848742033 | 0.5369755 | 1.41664E-05 |
| Decreasing, ↓ velocity | Activated CD4 T | RP11-573D15.2 | 4.817948172 | 0.6523456 | 1.52073E-05 |
| Decreasing, ↓ velocity | Activated CD4 T | SIPA1L3 | 4.816617681 | 0.6522441 | 1.5254E-05 |
| Increasing, ↑ velocity | Activated CD4 T | RPS21 | 4.789112694 | 0.5779985 | 1.62513E-05 |
| Increasing, ↑ velocity | Activated CD4 T | CWC22 | 4.78879827 | 0.6501161 | 1.6263E-05 |
| Max Parabolic | Mem B | AP000769.1 | 4.77523755 | 0.5195471 | 1.67789E-05 |
| Min Parabolic | NK | FBXL6 | 4.741740435 | 0.5290406 | 1.81242E-05 |
| Decreasing, ↓ velocity | XCL+ NKs | BASP1 | 4.736010959 | 0.7894152 | 1.83649E-05 |
| Decreasing, ↓ velocity | Activated CD4 T | PARP6 | 4.716626395 | 0.6445345 | 1.92032E-05 |
| Increasing, ↑ velocity | Activated CD4 T | CLPTM1L | 4.691242491 | 0.6425502 | 2.03591E-05 |
| Decreasing, constant | Activated CD4 T | ICAM2 | 4.682851526 | 0.5687815 | 2.07562E-05 |
| Increasing, ↑ velocity | NK | AC007950.2 | 4.677383817 | 0.5242029 | 2.10192E-05 |

| | | | | | |
|---|---|---|---|---|---|
| Max Parabolic | Activated CD4 T | VPS13B | 4.66171128 | 0.6402279 | 2.17916E-05 |
| Decreasing, constant | Activated CD4 T | IFI44L | 4.655968675 | 0.5664195 | 2.20816E-05 |
| Increasing, ↓ velocity | Mem B | RPS12 | 4.64722493 | 0.5100125 | 2.25307E-05 |
| Decreasing, ↑ velocity | Activated CD4 T | KIF2A | 4.620173098 | 0.5632553 | 2.39788E-05 |
| Decreasing, ↓ velocity | Naïve B | DSCC1 | 4.61839402 | 0.6033563 | 2.40772E-05 |
| Decreasing, ↑ velocity | Activated CD4 T | TXLNG | 4.617100309 | 0.5629827 | 2.4149E-05 |
| Min Parabolic | Naïve B | CYSTM1 | 4.614353158 | 0.6030353 | 2.43023E-05 |
| Decreasing, ↓ velocity | Naïve B | IFIT1 | 4.581863507 | 0.6004445 | 2.61901E-05 |
| Increasing, ↑ velocity | Activated CD4 T | MCOLN2 | 4.58079509 | 0.5597491 | 2.62546E-05 |
| Decreasing, constant | Activated CD4 T | SIAH2 | 4.56953922 | 0.558742 | 2.69439E-05 |
| Decreasing, ↓ velocity | Naïve B | DIRC2 | 4.553539882 | 0.5981722 | 2.7955E-05 |
| Increasing, ↑ velocity | Naïve B | HBEGF | 4.553430681 | 0.5981634 | 2.79621E-05 |
| Decreasing, ↓ velocity | XCL+ NKs | PPP1R3B | 4.53243984 | 0.774831 | 2.93468E-05 |
| Increasing, ↓ velocity | Activated CD4 T | RPS6 | 4.532295428 | 0.6298713 | 2.93565E-05 |
| Min Parabolic | XCL+ NKs | MNDA | 4.486940427 | 0.7714356 | 3.25881E-05 |
| Decreasing, ↓ velocity | XCL+ NKs | HNRNPCL1 | 4.443932927 | 0.7681792 | 3.59805E-05 |
| Increasing, ↓ velocity | Activated CD4 T | EEF2 | 4.431562412 | 0.6216041 | 3.70201E-05 |
| Max Parabolic | Activated CD4 T | TC2N | 4.423491415 | 0.6209338 | 3.77145E-05 |
| Decreasing, ↓ velocity | Naïve B | RP11-536K7.3 | 4.403889917 | 0.5859498 | 3.94557E-05 |
| Decreasing, ↓ velocity | Activated CD4 T | IGHV3-11 | 4.40093194 | 0.6190538 | 3.97254E-05 |
| Increasing, ↑ velocity | NK | BLNK | 4.395971153 | 0.5024582 | 4.01818E-05 |
| Decreasing, ↓ velocity | XCL+ NKs | TYMP | 4.387106524 | 0.7638051 | 4.10104E-05 |
| Decreasing, ↓ velocity | Activated CD4 T | BCYRN1 | 4.383012457 | 0.6175539 | 4.13988E-05 |
| Decreasing, ↑ velocity | Activated CD4 T | WIPF1 | 4.377957552 | 0.5412622 | 4.18835E-05 |
| Decreasing, ↓ velocity | Activated CD4 T | IGHV3-7 | 4.360075851 | 0.6156254 | 4.3644E-05 |
| Increasing, ↓ velocity | Activated CD4 T | TPT1 | 4.331912134 | 0.6132441 | 4.6568E-05 |
| Decreasing, constant | Naïve B | MX1 | 4.319941381 | 0.5046075 | 4.78695E-05 |
| Decreasing, ↓ velocity | XCL+ NKs | GOLGA8K | 4.305772758 | 0.7574007 | 4.94569E-05 |
| Decreasing, ↓ velocity | Activated CD4 T | IGLV3-27 | 4.305193493 | 0.6109714 | 4.9523E-05 |
| Decreasing, ↓ velocity | Activated CD4 T | NDUFB7 | 4.287548048 | 0.6094631 | 5.15765E-05 |
| Decreasing, ↑ velocity | Activated CD4 T | RAB14 | 4.240182331 | 0.5282898 | 5.75198E-05 |
| Max Parabolic | Activated CD4 T | HNRNPK | 4.234244372 | 0.6048712 | 5.83117E-05 |
| Decreasing, ↓ velocity | Activated CD4 T | BEND2 | 4.2257136 | 0.6041314 | 5.94684E-05 |
| Increasing, ↓ velocity | Activated CD4 T | RPS3A | 4.190545337 | 0.6010666 | 6.44844E-05 |
| Min Parabolic | Activated CD4 T | CCNI | 4.149177204 | 0.5974311 | 7.09288E-05 |
| Decreasing, constant | Activated CD4 T | SP110 | 4.129821378 | 0.5176509 | 7.41615E-05 |
| Increasing, ↑ velocity | Activated CD4 T | CALU | 4.118841566 | 0.5947441 | 7.60604E-05 |
| Decreasing, ↓ velocity | Activated CD4 T | S1PR3 | 4.115877379 | 0.5944806 | 7.65813E-05 |
| Increasing, ↓ velocity | Activated CD4 T | RPL27A | 4.108502406 | 0.5938242 | 7.78929E-05 |
| Min Parabolic | Naïve B | C6orf229 | 4.105171396 | 0.5604295 | 7.84926E-05 |

| | | | | | |
|---|---|---|---|---|---|
| Increasing, ↑ velocity | Activated CD4 T | SMG5 | 4.092344095 | 0.5923824 | 8.08455E-05 |
| Increasing, ↑ velocity | Activated CD4 T | TMOD1 | 4.079095907 | 0.512686 | 8.33497E-05 |
| Decreasing, ↓ velocity | Activated CD4 T | TMEM140 | 4.060847307 | 0.5895572 | 8.69266E-05 |
| Decreasing, ↓ velocity | Activated CD4 T | LAMP3 | 4.042452283 | 0.5878982 | 9.06876E-05 |
| Min Parabolic | Activated CD4 T | DCAF10 | 4.036613372 | 0.5873702 | 9.19151E-05 |
| Increasing, ↑ velocity | Activated CD4 T | SUCLG2 | 4.016240829 | 0.5064675 | 9.63295E-05 |
| Decreasing, ↓ velocity | Activated CD4 T | SUB1 | 4.00476555 | 0.5844783 | 9.89087E-05 |
| Decreasing, ↓ velocity | Naïve B | PPP1R32 | 3.93469484 | 0.5451663 | 0.000116227 |
| Decreasing, ↓ velocity | Activated CD4 T | IFI44 | 3.933838504 | 0.5779648 | 0.000116456 |
| Decreasing, ↓ velocity | Activated CD4 T | RP1-30M3.5 | 3.929746703 | 0.5775859 | 0.000117558 |
| Increasing, ↑ velocity | Activated CD4 T | C9orf85 | 3.892800209 | 0.5741495 | 0.000127997 |
| Decreasing, ↓ velocity | Activated CD4 T | RSAD2 | 3.891258791 | 0.5740056 | 0.000128452 |
| Increasing, ↑ velocity | Activated CD4 T | ARSA | 3.8732564 | 0.5723205 | 0.000133889 |
| Decreasing, ↓ velocity | Activated CD4 T | IGLV7-46 | 3.858190084 | 0.5709051 | 0.000138615 |
| Decreasing, ↓ velocity | Activated CD4 T | GEMIN6 | 3.851482794 | 0.5702735 | 0.000140772 |
| Decreasing, ↑ velocity | Activated CD4 T | ZC3H14 | 3.816531992 | 0.5669672 | 0.00015257 |
| Increasing, ↑ velocity | Activated CD4 T | TNFRSF13C | 3.766193709 | 0.5621606 | 0.000171319 |
| Decreasing, ↓ velocity | Activated CD4 T | IFIT3 | 3.752638858 | 0.5608571 | 0.000176751 |
| Min Parabolic | Activated CD4 T | RPGR | 3.730745842 | 0.5587437 | 0.000185889 |
| Decreasing, ↓ velocity | Activated CD4 T | FTCDNL1 | 3.712387476 | 0.5569637 | 0.000193916 |
| Increasing, ↑ velocity | Activated CD4 T | SLC4A1 | 3.710120649 | 0.5567434 | 0.00019493 |
| Min Parabolic | Activated CD4 T | PTAFR | 3.699309104 | 0.5556913 | 0.000199844 |
| Increasing, ↑ velocity | Activated CD4 T | AHSP | 3.656544729 | 0.551505 | 0.000220524 |
| Decreasing, ↓ velocity | Activated CD4 T | CCNE1 | 3.618961492 | 0.5477933 | 0.000240458 |
